# Berberine Alleviates Adiposity and Cardiac Dysfunction in Offspring Exposed to Gestational Diabetes Mellitus

**DOI:** 10.1101/2020.07.20.209395

**Authors:** Laura K. Cole, Li Chen, Genevieve C. Sparagna, Marilyne Vandel, Bo Xiang, Vernon W. Dolinsky, Grant M. Hatch

**Affiliations:** Diabetes Research Envisioned and Accomplished in Manitoba (DREAM) Theme, Children’s Hospital Research Institute of Manitoba, Department of Pharmacology & Therapeutics, Faculty of Health Sciences, University of Manitoba, Winnipeg, Canada.; Department of Pharmacology, College of Basic Medical Sciences, School of Nursing, Jilin University, Changchun, China; Department of Medicine, Division of Cardiology, University of Colorado Anschutz Medical Center, Aurora, USA.; Center for Research and Treatment of Atherosclerosis, University of Manitoba, Winnipeg, Canada.

**Author notes:** To whom correspondence should be addressed: Dr. Grant M. Hatch, 501C JBRC, 715 McDermot Avenue, Winnipeg, Manitoba, Canada, R3E 3P4.

**Keywords:** Berberine, Gestational Diabetes, Metabolism, Cardiolipin, Mitochondria

## Abstract

The most robust risk factor for type 2 diabetes in childhood is prior exposure to diabetes during gestation. Currently, there are few evidence-based strategies to attenuate the of risk of metabolic syndrome in offspring exposed to gestational diabetes mellitus (GDM). Berberine (BBR) is an isoquinoline alkaloid extracted from Chinese herbs and exhibits glucose lowering properties. It has been used safely for centuries in humans. Our objective was to determine whether BBR treatment improves health outcomes in the mouse offspring of GDM dams. Dams were fed either a lean low-fat diet (Lean, LF,10% kcal fat) or a GDM-inducing high-fat/high sucrose diet (GDM, HF, 45% kcal fat) prior to breeding and throughout pregnancy. The resulting Lean and GDM-exposed offspring were randomly assigned a LF, HF or HF diet containing BBR (160 mg/kg/d) for 12 weeks. We determined that BBR treatment significantly reduced body weight (∼20%), % body fat (∼40%) and gonadal fat pad mass (∼60%) compared to HF-fed GDM offspring. Furthermore, BBR treatment of HF-fed GDM offspring normalized insulin levels in the plasma and isolated pancreatic islets. Differences in food consumption did not contribute to altered body composition in BBR treated mice, as levels remained similar between experimental groups. Alternatively, BBR-treatment was associated with increased whole-body oxygen consumption (VO2), activity and heat production. Additionally, we determined that HF-fed GDM offspring developed a cardiomyopathy, characterized by increased isovolumetric contraction (∼150%, IVCT), relaxation time (∼70%, IVRT), elevated cardiac triglyceride (∼120%) and reduced mitochondrial function (30%, spare capacity) compared to LF fed Lean controls. BBR treatment normalized heart function, reduced triglyceride levels and maintained mitochondrial function. Our data supports BBR as a potential pharmacotherapeutic approach to improve health outcomes in individuals exposed to GDM.

**Key Points Summary:**

- Gestational diabetes mellitus is a common metabolic complication of pregnancy which is increasing worldwide due to the prevalence of obesity.
- It is known that individuals exposed to gestational diabetes have elevated risk of developing metabolic syndrome however there are few evidence-based strategies which provide protection.
- Berberine is a natural compound found in Chinese herbs which has been safely used for centuries to treat type 2 diabetes mellitus.
- We determined that berberine treatment of offspring exposed to gestational diabetes attenuated weight gain, reduced insulin levels and normalized both heart and pancreatic function.
- Our data supports berberine as a potential pharmacotherapeutic approach to improve health outcomes in individuals exposed to gestational diabetes.

## Introduction

Gestational diabetes mellitus (GDM) is defined as hyperglycemia caused by glucose intolerance during pregnancy and is the most common metabolic complication of gestation (Chiefari *et al*., 2017). The incidence of GDM continues to increase worldwide due to the prevalence of obesity and increasing maternal age. It is now clear that exposure to intrauterine hyperglycemia during GDM has long-term consequences for offspring health (Damm *et al*., 2016; Agarwal *et al*., 2018). Studies focused on the developmental origins of health and disease have determined that the offspring of women with GDM have significantly higher risk of developing obesity and prediabetes/diabetes compared to the background population (4 and 8-times respectively) (Clausen *et al*., 2008; Clausen *et al*., 2009; Nehring *et al*., 2013). The current treatment strategies for GDM (e.g. rigorous maternal lifestyle management, insulin injections and/or oral hypoglycemic drugs) lack universal compliance, due to practicality, difficulty learning the required skill set and/or elevated levels of anxiety (Chiefari *et al*., 2017). Furthermore, there are few evidence-based strategies to attenuate the risk of metabolic syndrome in offspring exposed to GDM. Thus, identifying effective therapeutic interventions for GDM exposed offspring are critical for improving public health.

Berberine (BBR) is an isoquinoline alkaloid enriched in plants of the genus *Berberis*. The root, stem, leaves and bark of these spiny shrubs (∼500 species) are widely considered to be the main natural source of BBR. BBR has been traditionally used to treat diabetes as an active constituent of the Chinese herbs huάng liάn (Coptis Root) and huang băi (Phellodendron Chinese) (Chang *et al*., 2015a; Ran *et al*., 2019). Numerous clinical trials have demonstrated that BBR (0.6-2.7 g/day for 2-3 months) is effective for treating hyperglycemia, and hyperlipidemia in patients with type 2 diabetes mellitus (Zhang *et al*., 2008; Lan *et al*., 2015). Zhang et al., determined that type 2 diabetes patients receiving BBR treatment (1 g/day) had significantly reduced fasting glucose (20%), postprandial glucose (25%) and triglyceride levels (35%) after 3 months (Zhang *et al*., 2008). Remarkably, the efficacy of BBR with respect to lowering fasting and postprandial glucose levels is similar to that of clinical hypoglycaemics (metformin, rosiglitazone) (Yin *et al*., 2008; Zhang *et al*., 2008; Lan *et al*., 2015). Clinical and preclinical studies have also demonstrated the beneficial effects of BBR on obesity (Lee *et al*., 2006; Sun *et al*., 2018), hepatic steatosis (Zhao *et al*., 2017) and cardiac dysfunction (Chang *et al*., 2015b; Dong *et al*., 2018; Feng *et al*., 2019). BBR is considered to have few side effects and no serious adverse reactions (Lan *et al*., 2015). Taken together, the low-cost, efficacy and tolerability of BBR have made this compound an attractive nutraceutical for the treatment of hyperglycemia, hyperlipidemia and type 2 diabetes. However, it remains unknown whether dietary BBR supplementation improves health outcomes in GDM exposed young adult offspring.

The present study represents the first *in vivo* GDM treatment strategy with BBR. We used a mouse model of diet-induced GDM to determine the effectiveness of post-weaning oral BBR supplementation on the metabolic health of high-fat fed offspring. Our research demonstrates that BBR treatment significantly reduced body weight (∼20%), % body fat (∼40%) and gonadal fat pad mass (∼60%) compared to HF-fed GDM offspring. Importantly, we determined for the first time that BBR treatment reduced adiposity in both male and female mice. BBR treatment of HF-fed GDM offspring also normalized insulin levels in the plasma and pancreatic β-cell function. Finally, we demonstrated that BBR treatment improved heart contractile function, prevented myocardial steatosis and maintained mitochondrial function in GDM exposed offspring.

## Methods

### Animal care and food preparation

This study was performed with approval of the University of Manitoba Animal Policy and Welfare Committee. All animals were maintained in an environmentally controlled facility (12 h light/dark cycle) with free access to food and water. We used a diet-induced GDM rodent model as previously described (Pereira *et al*., 2015; Brawerman *et al*., 2019). Briefly, female C57BL/6 mice (6 weeks old, obtained from University of Manitoba colony) were fed either a low-fat (10% kcal, Research Diets D12450B) or a high-fat/ sucrose diet (45% kcal, Research Diets D12451) for 6 weeks prior to mating with low-fat fed male C57BL/6 mouse (6-10 week old, obtained from the University of Manitoba colony) for 4 days. The diet assigned to the female mouse was continued during mating, pregnancy and suckling periods. The resulting litters were culled at a maximum of 8 to prevent nutritional deprivation of any offspring. At 3 weeks of age the male and female offspring were weaned and randomly assigned to one of the three diets 1) low fat (10% kcal) 2) high-fat/ sucrose (45% kcal) or 3) high fat/sucrose (45% kcal) diet supplemented with BBR (160 mg/kg/day) for a 12 week period (Fig. 1A).

**Figure 1:**
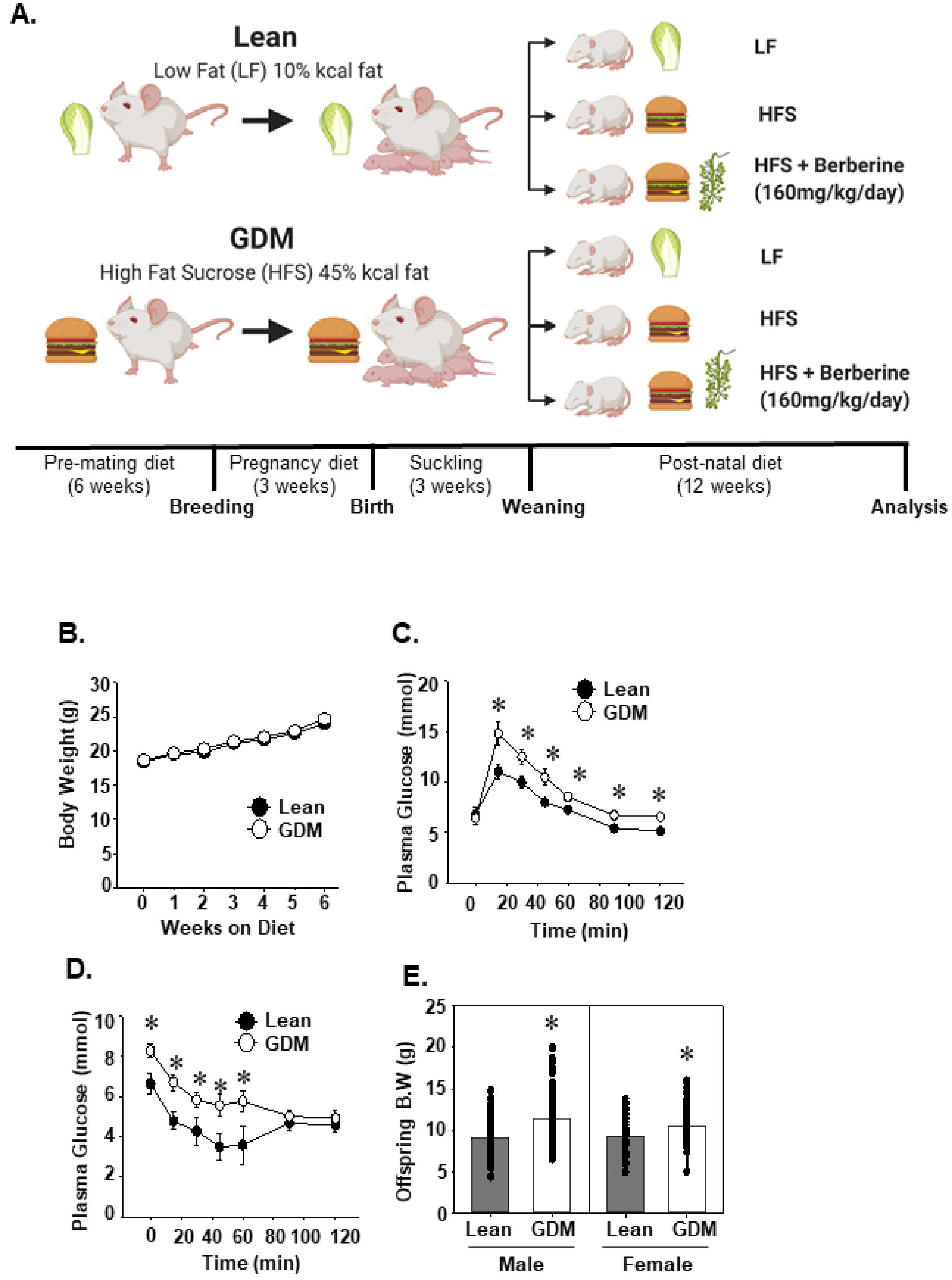
Characterization of lean and GDM dams. A, Lean and GDM dams were generated by feeding a low-fat or high-fat diet respectively for 6 weeks prior to mating. These diets were continued throughout pregnancy and the suckling period. B, Pre-mating maternal body weights (n=27-49). C, Maternal mid-gestation GTT and D, ITT (embryonic day 15) (n=6-12). E, Body weight of 3-week old offspring. Values are means ± SEMs. \**P*<0.05 compared with Lean mice.

Rodents will not eat traditional food pellets containing BBR due to the natural bitterness of this nutraceutical. As a result, previous studies have treated rodents with BBR by daily oral gavage or intraperitoneal injections (Lee *et al*., 2006; Sun *et al*., 2018). Alternatively, we prepared rodent food containing BBR by generating agar blocks using powered rodent diet purchased commercially from Research Diets (Low fat - D12450B, or high fat D12451). Briefly, agar 14.5 g/kg (Sigma) and the indicated LF or HF diet 435 g/kg were dissolved in heated water (1 kg) with or without BBR 1.45 g/kg (Sigma). The mixture was cooled at 4°C and cut into blocks which were freshly placed in animal hoppers every 2-3 days.

### In vivo assessment by echocardiography

Transthoracic echocardiography was performed as described (Cole *et al*., 2011) on mildly anesthetized animals with 1-1.5% isoflurane, 1L/min oxygen. Each animal was placed on a heated ECG platform to maintain body temperature and measure heart rate. A Vevo 2100 high resolution imaging system equipped with a 30-MHz transducer (Visual Sonics, Toronto) was used to visualize mouse hearts. The transducer was placed just below the sternum and angled toward the heart to obtain images for analysis of cardiac structures and function. Myocardial performance index was calculated using the equation (isovolumetric contraction time + isovolumetric relaxation time)/ejection time. Systemic blood pressure measurements were obtained using a tail cuff system (IITC Corp.). All assessments were performed by a technician blinded to the offspring gestational condition and diet.

### Metabolic parameters

Indirect calorimetry measurements were obtained using an Omnitech (Columbus, OH) animal monitoring system. Body lean and fat mass were determined by dual-energy x-ray absorptiometry scanning and expressed as a % of total body mass.

### Pancreatic islet isolation and hormone assays

Mice were anesthetized with pentobarbital (110 mg/kg) and the pancreas perfused with collagenase V via the common bile duct as previously described ((Agarwal *et al*., 2019). Islets were manually picked using a dissecting microscope (SZ61, Olympus) and incubated overnight at 37°C, 5% CO_2_ in 10% FBS, 1% P/S RPMI 1640. The following day, islets from each animal were divided (15 islets X 3 replicates). Insulin secretion experiments were performed by incubating each aliquot of islets at 37°C, 5% CO_2_ with Krebs-Ringer bicarbonate buffer (KRB) supplemented with 2.8 mM glucose for 1 h, followed by KRB containing 16.7 mM glucose for 30 mins, and KRB containing 16.7 mM glucose plus 3 mM KCL for 30 mins. The islets were then lysed (70% ethanol, 0.1 N HCl), dried down and resuspended in water. Total pancreatic islet insulin (islet), secreted insulin (ELISA, ALPCO) and total pancreatic islet glucagon (radioimmunoassay, Millipore) were quantitated and normalized to DNA content.

### Blood and tissue parameters

Following a 16 h fast the animals were anesthetized with 1-1.5% isoflurane and then euthanized by cervical dislocation. The plasma levels of insulin, and leptin were quantitated by ELISA (ALPCO). Commercially available kits were used to measure plasma non-esterified fatty acids, ketones, triglyceride (Wako Diagnositic), and alanine aminotransferase (Biotron Diagnostic). Molecular species of cardiolipin (CL) were quantitated from tissue homogenates by HPLC coupled to electrospray ionization mass spectrometry (Sparagna *et al*., 2005). The total CL was calculated from the sum of the eight most prominent CL species (1442, 1424, 1448, 1450, 1472, 1474, 1496, 1498) which were previously defined by fatty acid side chain composition (Sparagna *et al*., 2005).

### Mitochondrial analysis

Oxygen consumption rate (OCR) was measured from isolated mouse heart or liver mitochondria (2 µg) using an Agilent Seahorse Bioscience XF24 analyzer. The seahorse base media contained: 70 mM sucrose, 220 mM Mannitol, 5 mM MgCl_2_, 5 mM KH2PO4, 2 mM HEPES, 1 mM EGTA, 0.2 % BSA. Complex I was assayed from isolated mitochondria with media additionally containing 5 mM malate and 5 mM glutamate. Complex II was assayed from isolated mitochondria with media additionally containing 5 mM succinate and 2 µM rotenone. The following reagents were then added in the following order during the seahorse protocol: 8 mM ADP, 2.5 µM oligomycin, 8 µM FCCP, 4 µM antimycin A. State I was considered to be basal respiration prior to the addition of ADP. State III was considered to be ADP facilitated oxygen consumption (maximum respiration). State IV was considered oxygen consumption in the presence of oligomycin (ATP independent respiration). The addition of antimycin A inhibits all mitochondrial respiration therefore, any respiration in the presence of antimycin A was subtracted from all OCR values prior to calculating the average. The spare capacity was calculated using the following formula (State III -State I)/State I *100.

### RNA isolation and quantitative RT-PCR analysis

Total RNA was isolated from cardiac or liver tissue using an RNeasy® kit (Qiagen) and first-strand cDNA synthesis performed (2 µg total RNA) with SuperScript II (Invitrogen). PCR was performed using Eppendorf Realplex^2^ instrument with the gene specific primers (IDT) as indicated (Table 1). All mRNA levels were quantitated using a standard curve followed by normalization to the geometric mean (geomean) of Tata Box binding protein (TBP) and TFIIB (Vandesompele *et al*., 2002).

**Table 1:**
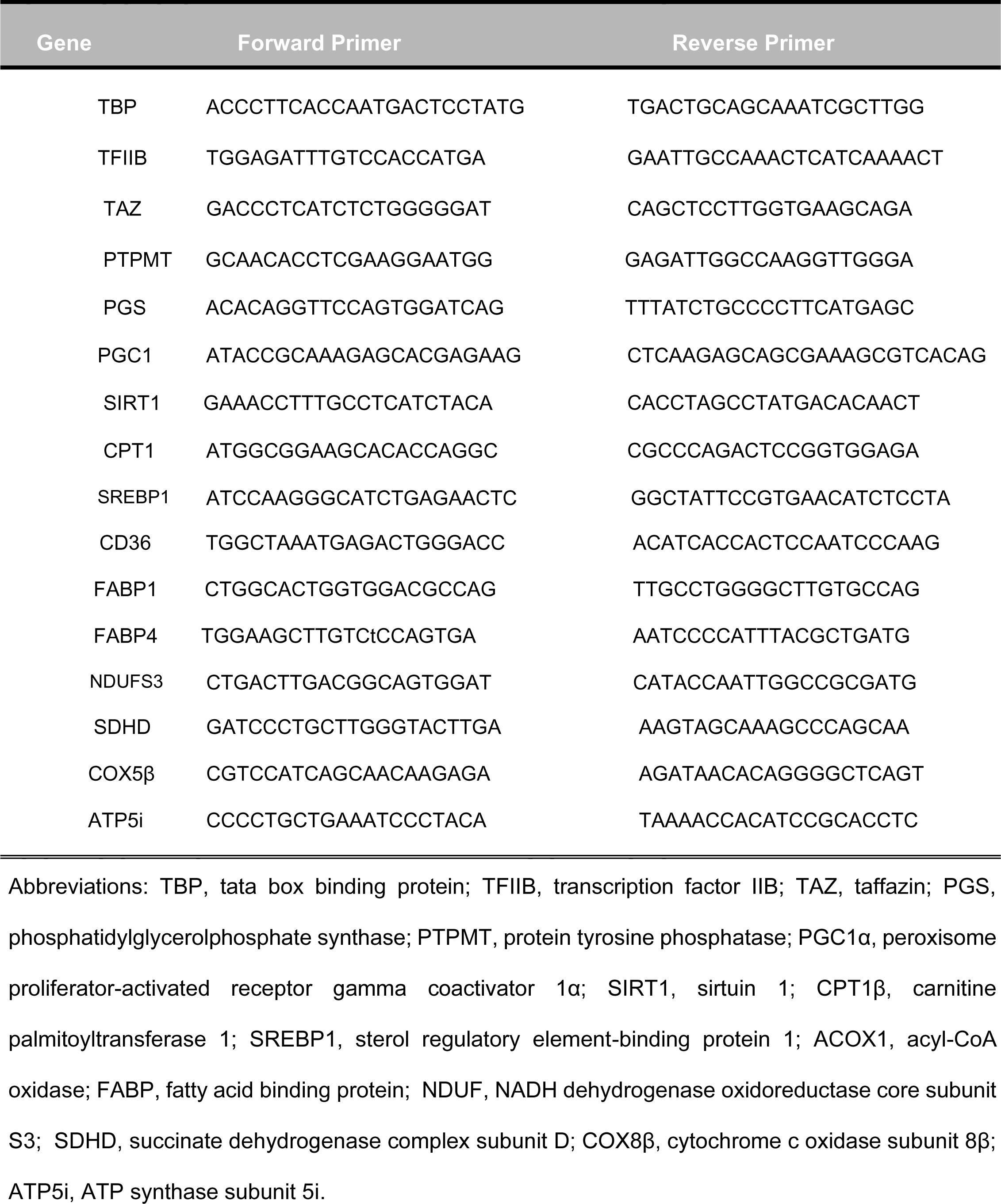
Primer sequences for quantitative PCR

### Statistical analysis

Data are expressed as means ± standard error of the mean (SEM). Comparisons between Lean and GDM offspring groups fed the same experimental diet was performed by unpaired two-tailed Student’s t-test. Comparisons between experimental diets (LF, HF and HFB) of the same gestational group (Lean and GDM) were determined using analysis of variance using Student-Newman-Keuls post-hoc analysis. A probability value of <0.05 was considered significant.

## Results

### Maternal characteristics

To generate Lean and GDM groups we fed C57BL/6 dams either a LF (10% kcal) or a HF diet (45% kcal) respectively for 6-weeks prior to mating (Fig. 1A). During this feeding period there was no difference in body weight between the dietary groups (Fig. 1B). To confirm *in utero* exposure of GDM fetuses to hyperglycemia we performed glucose tolerance tests (GTT) during the second trimester of pregnancy (embryonic day 15). Consistent with previous studies, at mid-gestation the GDM dams exhibited impaired glucose tolerance with significant increases in plasma glucose levels at multiple GTT time points compared to the Lean dam controls (Fig. 1C) (Pereira *et al*., 2015; Brawerman *et al*., 2019). Similarly, pregnant GDM dams had impaired insulin sensitivity compared to the Lean Dams (Fig. 1D). Despite the lack of change in body weight during the pre-mating period, GDM dams weighted significantly more postnatally (3-4 weeks post-partum) than the Lean controls (Table 2). The increased body weight was accompanied by elevated mass of gonadal white adipose tissue in the GDM dams (Table 2).

**Table 2.** Maternal characteristics from post-natal Lean and GDM dams. All metabolic characteristics of dams were determined following the weaning of offspring at 3 weeks of age. Values are means ± SEMs (n=14-49). Tissue weights are expressed per g of body weight. *\*P<0.05* compared with Lean dams.

|  | LEAN Dams | GDM Dams |
| --- | --- | --- |
| Weights (g) | 25.92±0.56 | 29.86±0.72* |
| **Plasma** |  |  |
| Triglyceride (mg/dL) | 6.38±0.85 | 6.46±0.54 |
| NEFA (mmol/L) | 0.45±0.05 | 0.35±0.02 |
| **Tissue** |  |  |
| Liver (g/g) | 0.048±0.003 | 0.033±0.001* |
| Gonadal Adipose (g/g) | 0.021±0.003 | 0.036±0.002* |
| Heart (g/g) | 0.0058±0.00002 | 0.0046±0.0002* |

### BBR treatment attenuated GDM-induced obesity in male and female offspring

At 3-weeks of age, male and female offspring were weaned and randomly assigned a LF (10% kcal), HF (45% kcal) or HF diet supplemented with BBR (HFB, 160 mg/kg/day) for a 12-week period (Fig. 2A). The body weights of the offspring were monitored weekly during the dietary treatments (Fig. 2A-D). Consistent with previous reports, GDM-exposure significantly increased the weight of both male and female offspring (3-weeks of age, Fig. 2E) (Brawerman *et al*., 2019). The increase in body weight for male GDM-exposed offspring persisted until the end of the 12-week period due to elevated adiposity (Fig. 2E and G, Fig 3A-D). As expected, BBR significantly decreased the body weight of offspring exposed to GDM as a result of significant reduction in whole body fat content (Fig. 2G) with smaller fad pads (Fig. 3A-D) and reduced plasma leptin levels (Table 3) (Lee *et al*., 2006; Gomes *et al*., 2012). Dietary BBR also reduced body weight and adiposity of male offspring within the Lean gestational group (Fig. 2E ad G). The percentage of lean mass was greater in the HFB fed groups compared to both the LF and HF fed animals regardless of the gestational condition (Fig. 2I).

**Figure 2:**
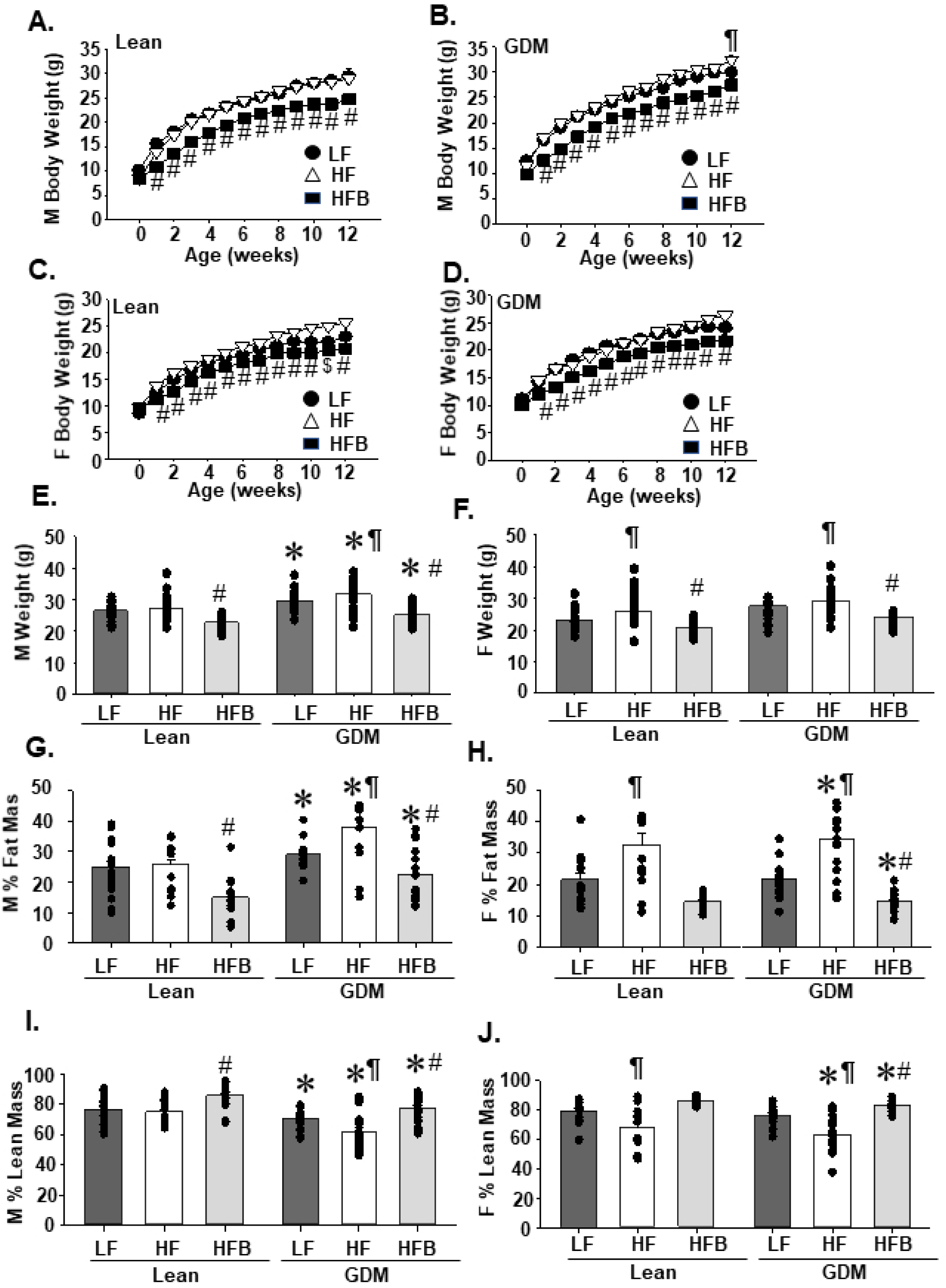
The body weight and fat mass of offspring is reduced by berberine treatment. Growth curves for male offspring from Lean A, and GDM B, dams (n=23-35). Growth curves for female offspring of C, Lean and D, GDM dams (n=17-26). Body weights for E, male and F, female offspring of lean and GDM dams following 12 weeks on the low-fat (LF), high-fat (HF) or high-fat berberine (HFB) diet. The % fat and lean body mass were calculated for male G, I, and female H, J, offspring, respectively. Values are means ± SEMs. \**P*<0.05 compared with offspring from lean dams. ¶ *P*<0.05 compared to LF and HFB fed offspring from the same gestational condition. #*P*< compared to LF and HF fed offspring from the same gestational condition.

**Figure 3:**
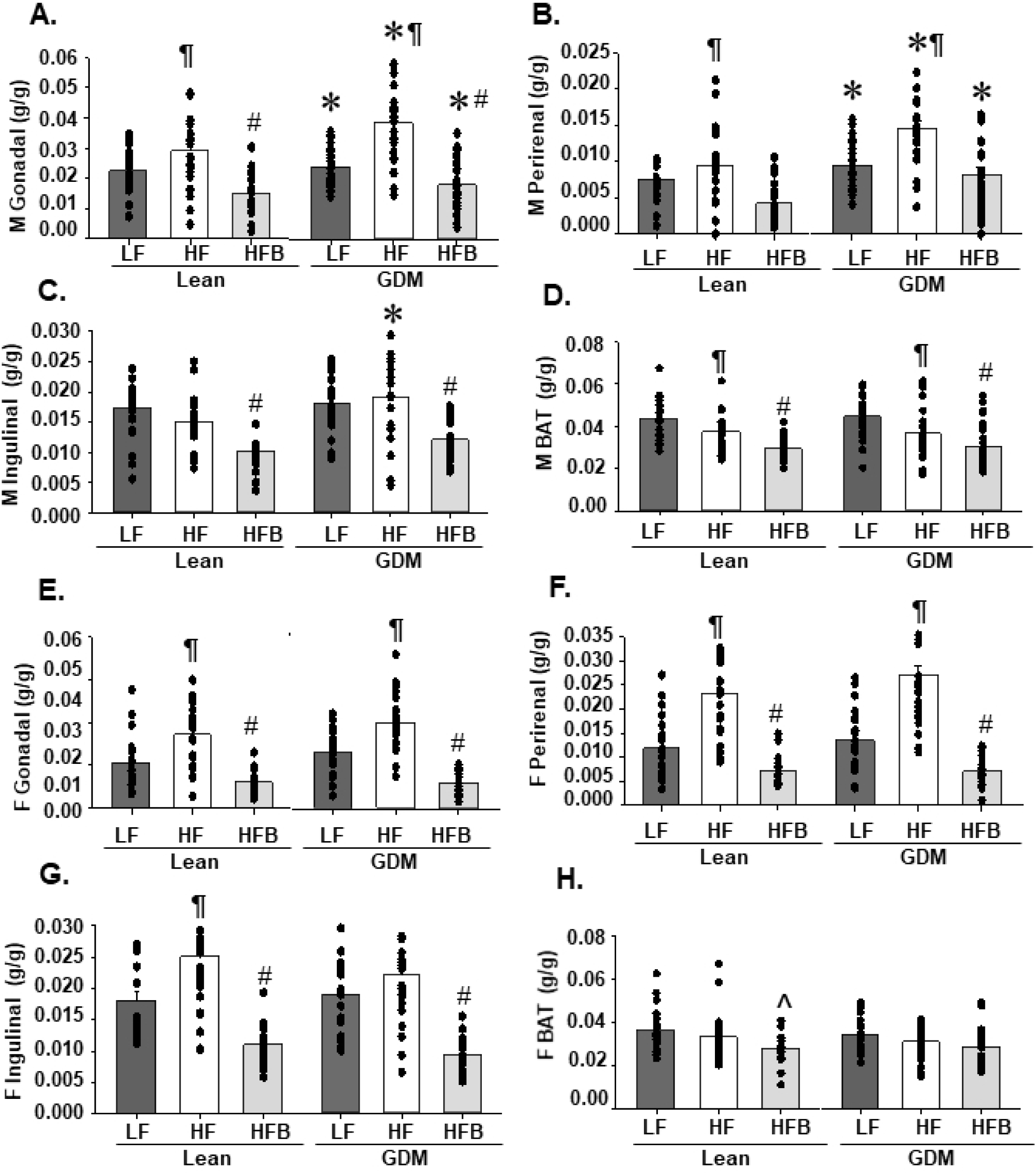
Fat pad mass is reduced by berberine treatment in offspring of lean and GDM dams. The mass of A, gonadal, B, perirenal, C, inguinal and D, BAT for male offspring of lean and GDM dams following 12 weeks of the indicated diet. The mass of E, gonadal, F, perirenal, G, inguinal and H, BAT for female offspring of lean and GDM dams following 12 weeks of the indicated diet. Values are means ± SEMs. \**P*<0.05 compared with offspring from lean dams. ¶ *P*<0.05 compared to LF and HFB fed offspring from the same gestational condition. ^#^*P*< compared to LF and HF fed offspring from the same gestational condition. ^*P*<0.05 compared to LF fed offspring from the same gestational condition.

**Table 3.** Characteristics of male offspring. All measurements were obtained following 12-weeks of the indicated diet (LF, HF, HFB). Values are means ± SEMs (n=5-15). \**P*<0.05 compared with offspring from lean dams. ¶*P*<0.05 compared to LF and HFB fed offspring from the same gestational condition. ^#^*P*< compared to LF and HF fed offspring from the same gestational condition. ^$^*P*<0.05 compared to HF fed offspring from the same gestational condition. NEFA; non-esterified fatty acids, ALT, alanine aminotransferase.

|  | LEAN<br>LF | LEAN<br>HF | LEAN<br>HFB | GDM<br>LF | GDM<br>HF | GDM<br>HFB |
| --- | --- | --- | --- | --- | --- | --- |
| **Consumption (24 h)** |  |  |  |  |  |  |
| Water (g/g) | 0.009±0.002 | 0.008±0.001 | 0.013±0.002 | 0.008±0.002 | 0.009±0.002 | 0.017±0.003<sup>#</sup> |
| Food (g/g) | 0.56±0.06 | 0.41±0.04 | 0.54±0.03<sup>§</sup> | 0.49±0.05 | 0.40±0.07 | 0.48±0.04 |
| Food (kcal/g) | 2.13±0.24 | 1.94±0.20 | 2.54±0.14 | 1.88±0.21 | 1.90±0.32 | 2.24±0.20 |
| **Plasma** |  |  |  |  |  |  |
| Glucose (mmol/L) | 5.36±0.31 | 5.26±0.25 | 5.88±0.24 | 5.47±0.35 | 4.83±0.16 | 6.42±0.28<sup>#</sup> |
| Insulin (ng/mL) | 0.064±0.01 | 0.093±0.02 | 0.044±0.01 | 0.210±0.04<sup>*</sup> | 0.244±0.06<sup>*</sup> | 0.105±0.01<sup>*#</sup> |
| Leptin (ng/mL) | 1.36±0.38 | 2.65±0.85 | 0.37±0.14<sup>§</sup> | 2.90±0.75 | 8.07±1.60<sup>*¶</sup> | 0.78±0.20 |
| NEFA (mmol/L) | 0.40±0.03 | 0.46±0.04 | 0.30±0.03<sup>#</sup> | 0.39±0.02 | 0.36±0.04 | 0.34±0.03 |
| Triglycerides (mg/dL) | 19.98±1.33 | 27.51±3.22 | 17.90±2.44<sup>§</sup> | 22.62±2.48 | 29.62±3.55 | 29.66±4.01<sup>*</sup> |
| Ketones (mmol/L) | 0.44±0.07 | 0.51±0.05 | 0.69±0.17 | 0.70±0.09<sup>*</sup> | 0.88±0.16<sup>*</sup> | 0.53±0.06 |
| ALT (IU/L) | 35.8±2.7 | 28.9±3.7 | 30.9±3.0 | 40.5±4.9 | 46.7±8.4<sup>*</sup> | 37.7±4.9 |

In female offspring, GDM exposure lacked any significant effect on body weight following the 12-week post-natal feeding period (Fig. 2F). A similar finding was previously determined in females using the model of obesity-induced GDM in rats (Pereira *et al*., 2015). Alternatively, the effect of HF diet was more pronounced in female offspring with significant increases in weight and adiposity within both the Lean and GDM exposed groups (Fig. 2F and H). The HF fed Lean and GDM offspring had elevated fat pad mass (Fig. 2H, Fig. 3E-H) and significantly increased plasma leptin levels (Table 4). BBR treatment prevented the HF diet mediated increases in body weight, fat pad mass and plasma leptin levels observed in female offspring (Fig. 2 F, H and J, Fig. 3E-H, Table 4), Together these findings indicate that BBR treatment reduces body weight and attenuated adiposity in male and female offspring exposed to GDM exposed and fed a HF diet.

**Table 4.** Characteristics of female offspring. All measurements were obtained following 12-weeks of the indicated diet (LF, HF, HFB). Values are means ± SEMs (n=3-15). \**P*<0.05 compared with offspring from lean dams. ¶*P*<0.05 compared to LF and HFB fed offspring from the same gestational condition. ^#^*P*< compared to LF and HF fed offspring from the same gestational condition. NEFA; non-esterified fatty acids, ALT, alanine aminotransferase.

|  | LEAN<br>LF | LEAN<br>HF | LEAN<br>HFB | GDM<br>LF | GDM<br>HF | GDM<br>HFB |
| --- | --- | --- | --- | --- | --- | --- |
| **Consumption (24 h)** |  |  |  |  |  |  |
| Water (g/g) | 0.013±0.002 | 0.005±0.002 | 0.014±0.005 | 0.007±0.002 | 0.009±0.004 | 0.020±0.004<sup>#</sup> |
| Food (g/g) | 0.68±0.07 | 0.60±0.06 | 0.73±0.03 | 0.68±0.05 | 0.59±0.05 | 0.60±0.04 |
| Food (kcal/g) | 2.61±0.25 | 2.83±0.29 | 3.41±0.13 | 2.61±0.18 | 2.79±0.24 | 2.98±0.24 |
| **Plasma** |  |  |  |  |  |  |
| Glucose (mmol/L) | 4.92±0.44 | 5.06±0.30 | 6.15±0.43 | 5.00±0.38 | 5.26±0.22 | 5.66±0.25 |
| Insulin (ng/mL) | 0.052±0.01 | 0.112±0.02<sup>¶</sup> | 0.034±0.01 | 0.075±0.02 | 0.167±0.04<sup>¶</sup> | 0.053±0.01 |
| Leptin (ng/mL) | 1.19±0.23 | 2.49±0.47<sup>¶</sup> | 0.72±0.16 | 1.97±0.73 | 6.20±1.03<sup>*¶</sup> | 1.06±0.33 |
| NEFA (mmol/L) | 0.30±0.03 | 0.29±0.02 | 0.26±0.02 | 0.30±0.03 | 0.26±0.03 | 0.25±0.03 |
| Triglycerides (mg/dL) | 9.94±1.45 | 16.17±2.10 | 16.14±1.49 | 12.11±2.79 | 14.09±1.22 | 17.73±1.64 |
| Ketones (mmol/L) | 0.71±0.03 | 0.65±0.06 | 0.88±0.10 | 0.86±0.09 | 1.15±0.21<sup>*</sup> | 0.80±0.09 |
| ALT (IU/L) | 39.9±2.0 | 23.7±5.2 | 29.3±4.4 | 39.9±3.3 | 26.6±6.0 | 22.5±2.7 |

### BBR promotes hypermetabolism in male and female GDM offspring

The BBR-mediated reduction in adiposity could not be attributed to decreases in food or water intake (Table 3 and Table 4). Therefore, to further assess the protective effect of dietary BBR we performed indirect calorimetry. In the male offspring, we determined that GDM exposure significantly reduced oxygen consumption, heat production and activity levels in the LF and/or HF fed offspring (Fig. 4A-F). Dietary BBR prevented GDM-induced reductions to whole-body metabolism by promoting oxygen consumption, heat production and activity (Fig. 4A-F). The respiratory exchange ratio was higher in the LF fed groups regardless of gestational condition due to the expected shift toward increased fat metabolism with HF diets (Fig. 4G and H). In the female offspring, we did not detect any significant effect of GDM exposure or HF diet on indirect calorimetry measurements (Fig. 5A-H). However, BBR treatment significantly increased both oxygen consumption and heat production in GDM exposed female offspring (Fig. 5A-F). Thus, our data indicates that BBR treatment attenuates obesity in both male and female GDM exposed offspring in part by elevating whole-body metabolism.

**Figure 4:**
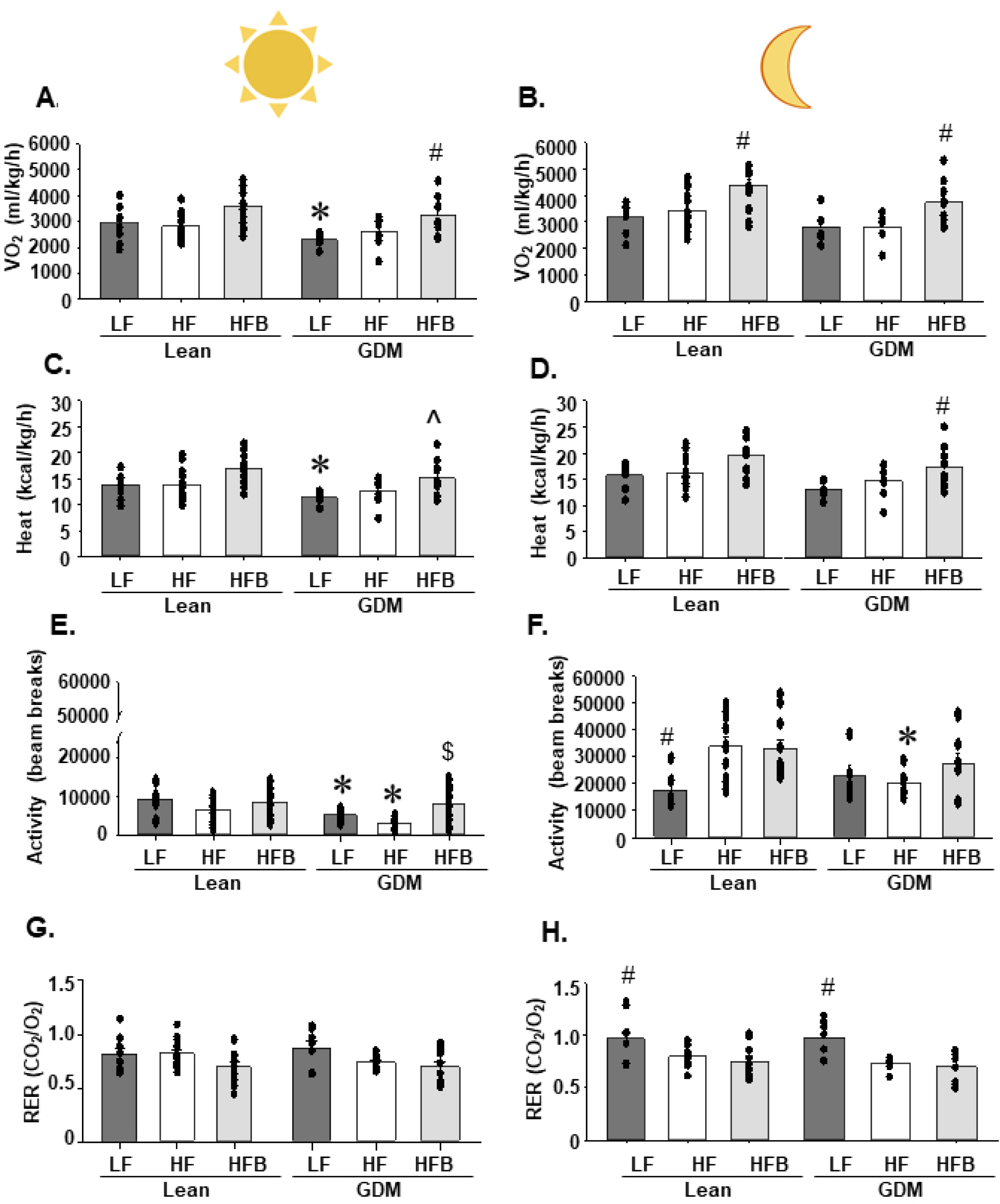
Berberine elevates metabolism and activity in male GDM offspring. Indirect calorimetry was performed for 24 h to measure A, B, oxygen consumption, C, D, heat production, E, F, activity and G,H the respiratory exchange ratio (RER) during the day and night, respectively. All measurements were performed following 12 weeks of the indicated diet. Values are means ± SEMs. \**P*<0.05 compared with offspring from lean dams. ¶ *P*<0.05 compared to LF and HFB fed offspring from the same gestational condition. ^#^*P*< compared to LF and HF fed offspring from the same gestational condition. ^*P*<0.05 compared to LF fed offspring from the same gestational condition. ^$^*P*<0.05 compared to HF fed offspring from the same gestational condition.

**Figure 5:**
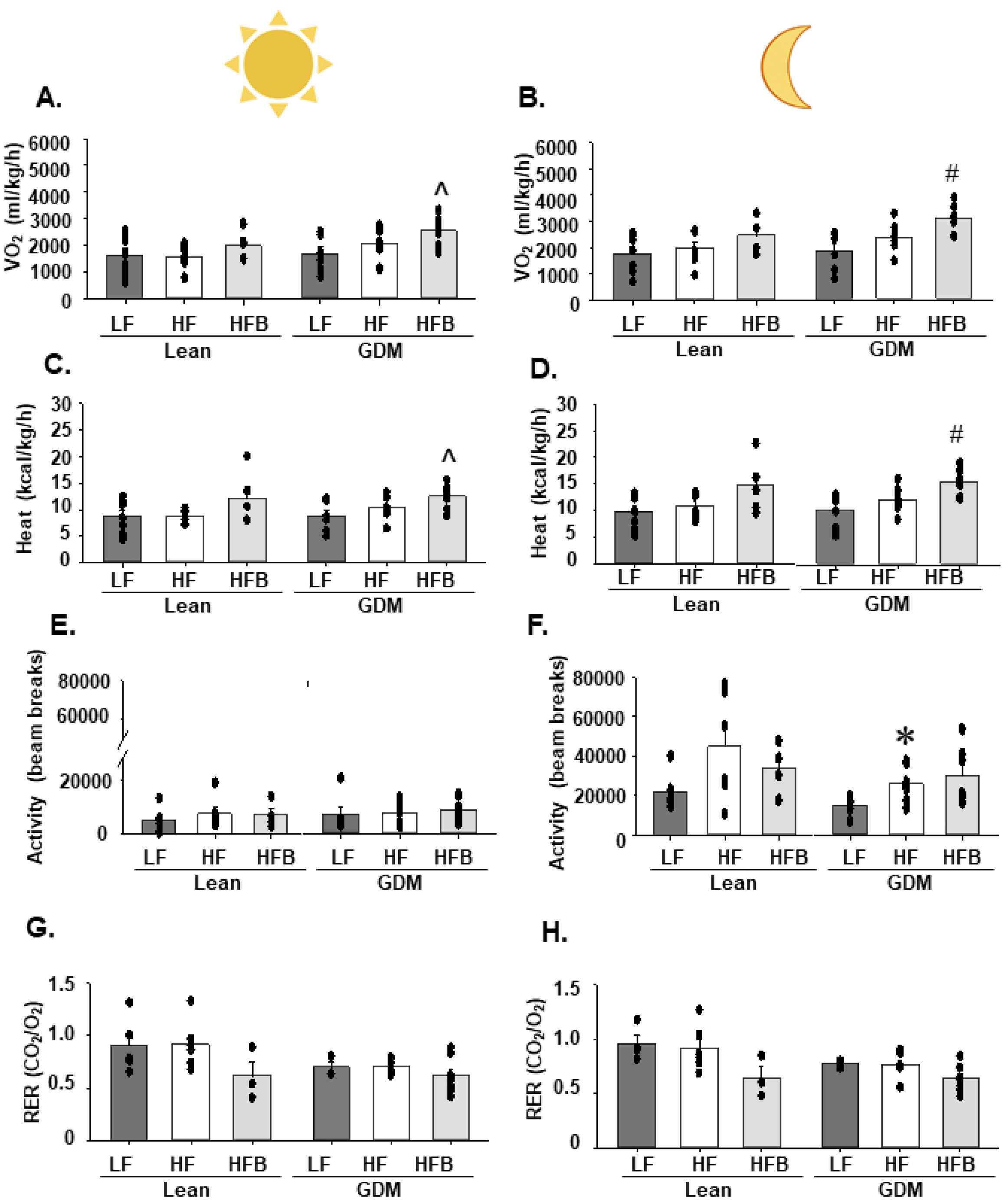
Berberine elevates metabolism in female GDM offspring. Indirect calorimetry was performed for 24 h to measure A, B, oxygen consumption, C,D, heat production, E, F, activity and G,H the respiratory exchange ratio (RER) during the day and night, respectively. All measurements were performed following 12 weeks of the indicated diet. Values are means ± SEMs. \**P*<0.05 compared with offspring from lean dams. ¶ *P*<0.05 compared to LF and HFB fed offspring from the same gestational condition. ^#^*P*< compared to LF and HF fed offspring from the same gestational condition. ^*P*<0.05 compared to LF fed offspring from the same gestational condition.

### BBR treatment protected pancreatic islets from dysfunction associated with GDM exposure in male offspring

BBR treatment has previously been shown to improve glucose homeostasis in clinical and preclinical models of diabetes (Lee *et al*., 2006; Yin *et al*., 2008; Zhang *et al*., 2008; Lan *et al*., 2015). In our experiments, GDM exposure did not promote hyperglycemia or widespread reductions in glucose and insulin tolerance (Tables 3 and 4, Fig. 6A-H). Therefore, our ability to assess the protective effects of BBR on these parameters was limited. Alternatively, GDM exposure promoted hyperinsulinemia in male offspring which was normalized to within control values (LF Lean group) with BBR (Table 3).

**Figure 6:**
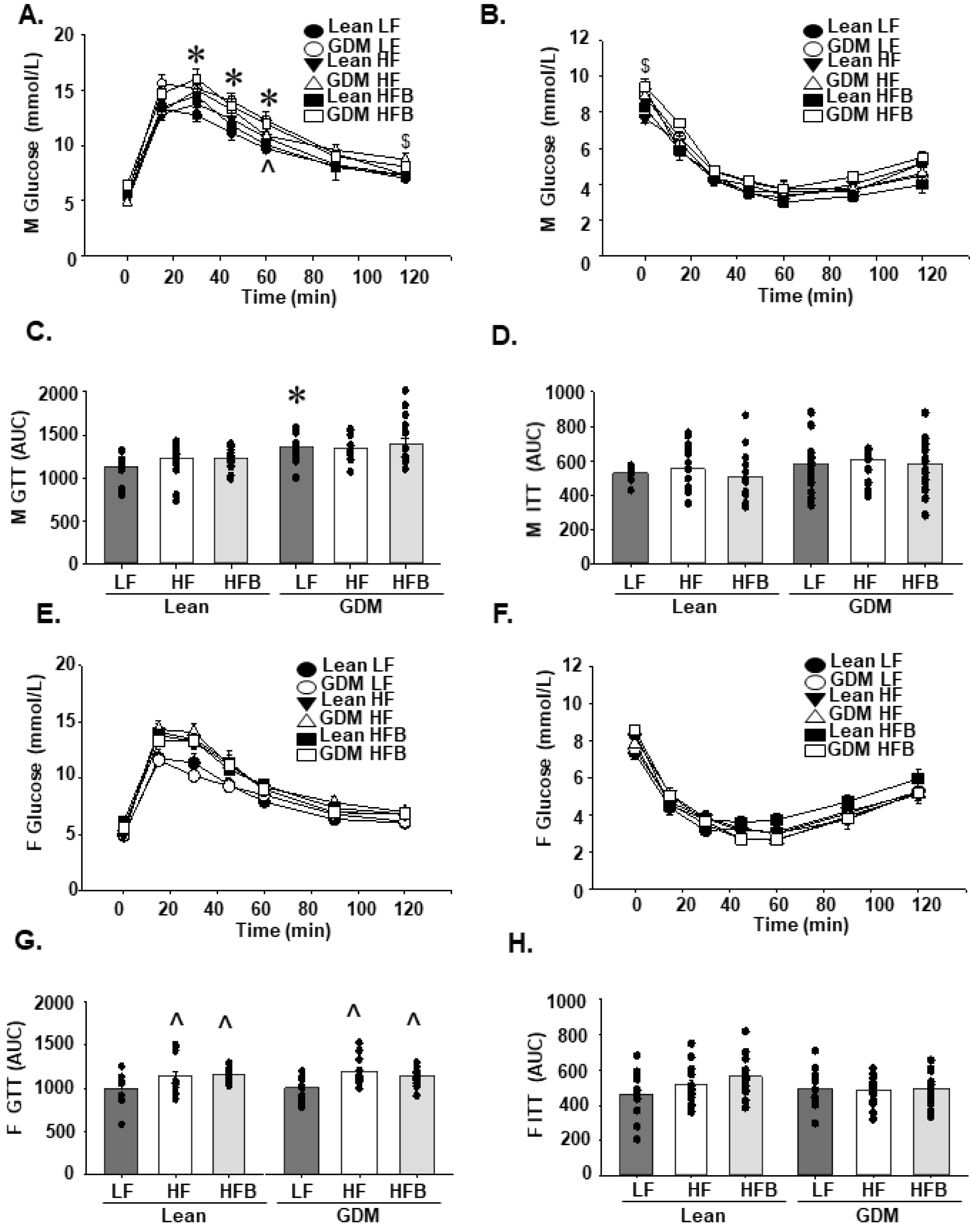
Offspring glucose homeostasis. A, GTT for male offspring. B, ITT for male offspring. C, GTT area under the curve for male offspring. D, ITT area under the curve for male offspring. E, GTT for female offspring. F, ITT for female offspring. G, GTT area under the curve for female offspring. H, ITT area under the curve for female offspring. All measurements were obtained following 12 weeks of the indicated diet (LF, HF, HFB). Values are means ± SEMs. \**P*<0.05 compared with offspring from lean dams. ¶*P*<0.05 compared to LF and HFB fed offspring from the same gestational condition. ^#^*P*< compared to LF and HF fed offspring from the same gestational condition. ^*P*<0.05 compared to LF fed offspring from the same gestational condition.

In an attempt to understand the hypoinsulinemic effect of BBR, we evaluated the function of pancreatic islets isolated from male offspring. We determined that islet insulin content was elevated by HF diet but was not further altered by GDM. Indicating that GDM-induced hyperinsulinemia was not due to enhanced hormone biosynthesis (Fig. 7A and B). To assess the secretory function of pancreatic β-cells, we measured insulin secretion rates under both low (2.8 mM) and high (16.7 mM) glucose conditions (Fig. 7C). Interestingly, the combined effect of GDM exposure and HF diet significantly increased basal (low-glucose) insulin secretion (Fig. 7C). With high glucose conditions, insulin secretion rates were similar between experimental groups (Fig. 7C), therefore the GDM HF fed group had reduced glucose-stimulated insulin secretion (GSIS) (Fig. 7D). A similar reduction in GSIS was detected for the LF fed GDM exposed animals (Fig. 7D). The dysfunction in basal insulin secretion was not a result of defective secretion machinery since the rates of insulin secretion in the presence of depolarizing conditions (KCL) was unchanged between the various groups of male offspring (Fig. 7E). Interestingly, basal insulin hypersecretion is linked to hyperinsulinemia and the progression to type 2 diabetic (Thomas *et al*., 2019). Dietary treatment with BBR normalized basal insulin secretion rates and GSIS in HF fed male offspring (Fig. 7A, C and D). Together these data indicate that BBR treatment protects islets from dysfunction associated with insulin hypersecretion in GDM exposed offspring.

**Figure 7:**
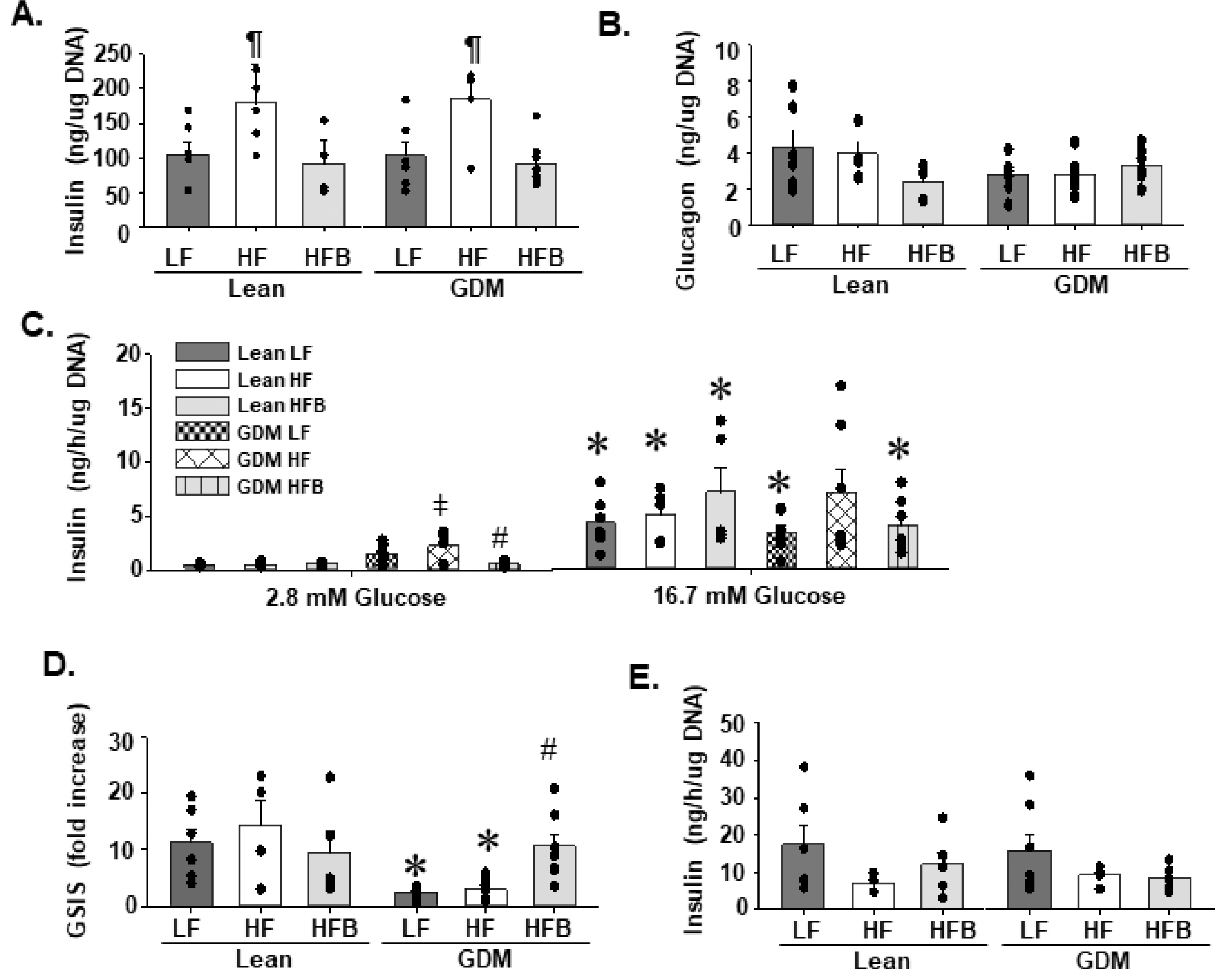
Berberine treatment prevented elevated insulin levels in pancreatic islets from male GDM offspring. A, Total insulin content in isolated islets. B, total glucagon content in isolated islets. C, Insulin secretion rates from isolated islets. D, Glucose-stimulated insulin secretion (GSIS) from isolated islets. E, insulin secretion rate in response to high glucose (16.7 mM) and KCl (3 mM). All measurements were obtained from male offspring following 12 weeks of the indicated diet (LF, HF, HFB). Values are means ± SEMs. \**P*<0.05 compared with offspring from lean dams. ¶*P*<0.05 compared to LF and HFB fed offspring from the same gestational condition. ^#^*P*< compared to LF and HF fed offspring from the same gestational condition. ^*P*<0.05 compared to LF fed offspring from the same gestational condition. ^ǂ^*P*<0.05 compared to Lean LF, HF, HFB and GDM HFB.

### Minimal changes to hepatic parameters in response to BBR treatment

There are numerous studies on the beneficial effects of BBR on the prevention of hepatic steatosis (Yuan *et al*., 2015; Zhao *et al*., 2017; Sun *et al*., 2018). Since, GDM has previously been shown to promote hepatic steatosis (Pereira *et al*., 2015; Brawerman *et al*., 2019), we investigated whether BBR would be beneficial in mice. We determined that, GDM exposure did not promote accumulation of TG in the liver. Consistent with this result GDM exposure did not increase plasma lipid levels (TG, NEFA) or alter the expression hepatic of FA metabolism enzymes (Table 5). While significant modification of hepatic gene expression occurred in response to HF diet for both FA uptake (FATBP1, FATBP4) and FAO (PGC1a, SIRT1), these levels were not further augmented by GDM exposure (Table 5). When BBR was supplemented in the diet, a significant reduction in hepatic TG levels was limited to the Lean gestational group (HFB Lean, Fig. 8A). Consistent with previous BBR studies, this result was mirrored by significant reductions in plasma lipid levels (TG, NEFA) (Table 3) and altered hepatic gene expression (increased FAO and reduced FA uptake) in the HFB Lean offspring (Table 5) (Yuan *et al*., 2015; Zhao *et al*., 2017).

**Figure 8:**
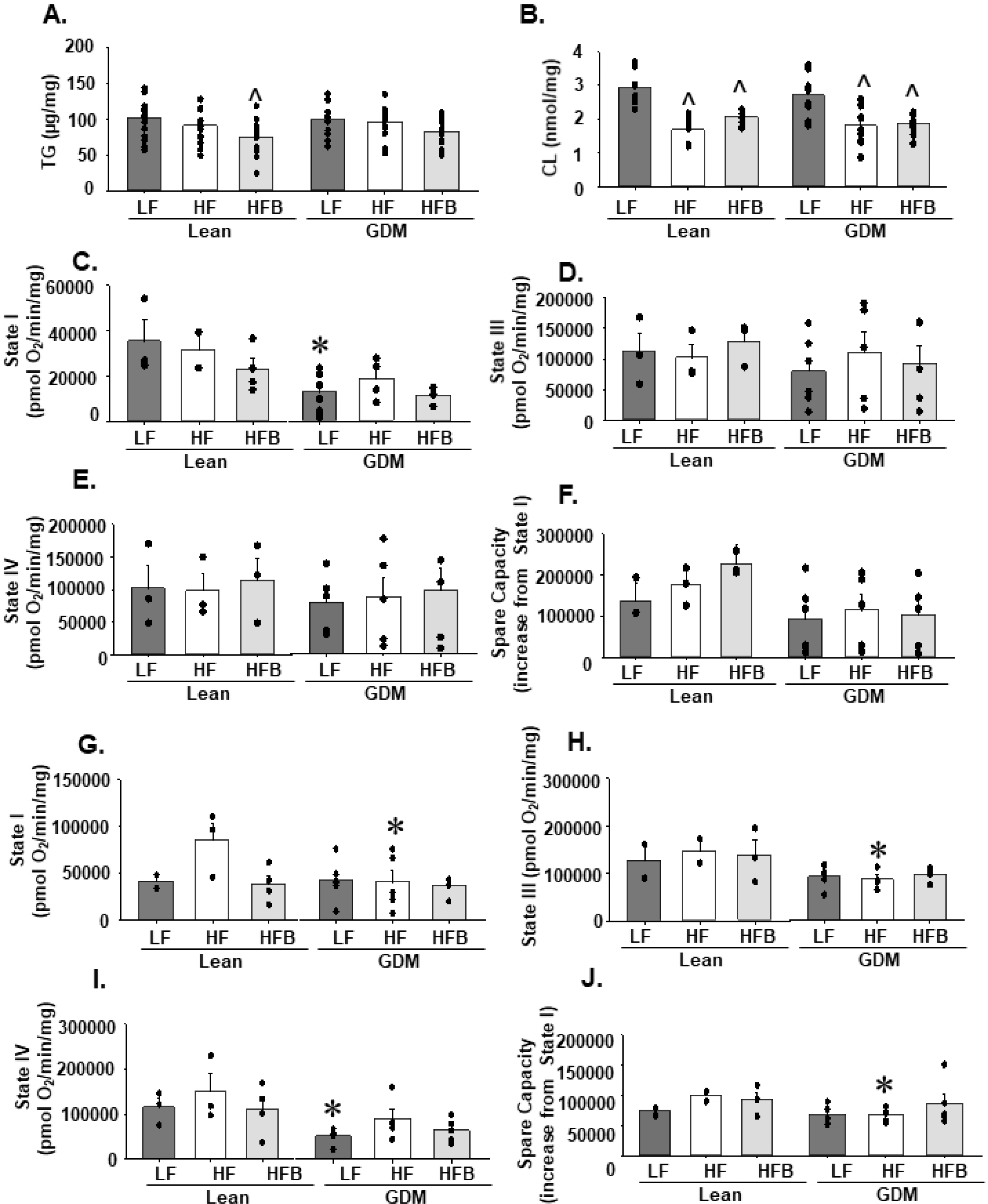
Hepatic mitochondrial function was not altered by berberine treatment. A, Hepatic triglyceride (TG) content and B, total cardiolipin content of male offspring. The oxygen consumption rates (OCR) were measured from hepatic mitochondria isolated from male offspring of Lean and GDM dams fed the indicated diet for 12 weeks. C, Oxygen consumption was measured for Complex I in respiratory state I (basal respiration, substrate in the absence of ADP), D, state III (maximum respiration) and, E state IV (oligomycin (ATP)-independent respiration). F, Spare capacity was calculated using the following formula (state III – state I) / state I *100%). Oxygen consumption was also measured for Complex II in G, respiratory state I (basal respiration, substrate in the absence of ADP), H, state III (maximum respiration, I, state IV (oligomycin (ATP)-independent respiration) and J, spare capacity. All data was normalized to mitochondrial protein content. Values are means ± SEMs. \**P*<0.05 compared with offspring from lean dams. ^*P*<0.05 compared to LF fed offspring from the same gestational condition.

**Table 5:**
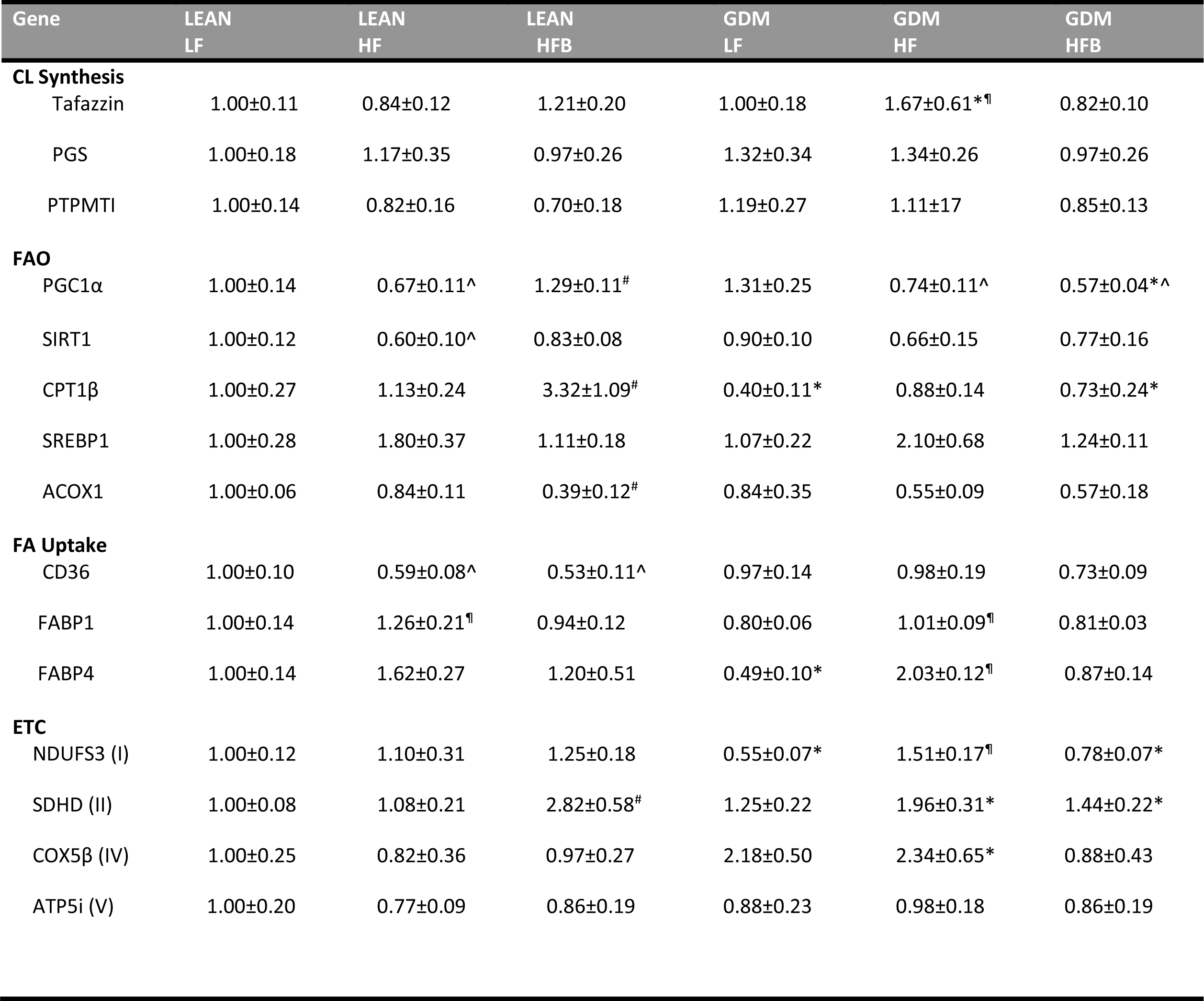
PCR analysis of liver. mRNAs were measured in the liver of offspring following 12-weeks of the indicated diet (LF, HF, HFB). Values are means ±SEMs (n=7). \**P*<0.05 compared with offspring from lean dams. ¶*P*<0.05 compared to LF and HFB fed offspring from the same gestational condition. ^#^*P*< compared to LF and HF fed offspring from the same gestational condition. ^*P*<0.05 compared to LF fed offspring from the same gestational condition. ^$^*P*<0.05 compared to HF fed offspring from the same gestational condition. The abbreviations for the mRNAs are: TAZ, taffazin; PGS, phosphatidylglycerolphosphate synthase; PTPMT, protein tyrosine phosphatase; PGC1α, peroxisome proliferator-activated receptor gamma coactivator 1α; SIRT1, sirtuin 1; CPT1β, carnitine palmitoyltransferase 1; SREBP1, sterol regulatory element-binding protein 1; ACOX1, acyl-CoA oxidase; FABP, fatty acid binding protein; NDUF, NADH dehydrogenase oxidoreductase core subunit S3; SDHD, succinate dehydrogenase complex subunit D; COX8β, cytochrome c oxidase subunit 8β; ATP5i, ATP synthase subunit 5i.

GDM-exposure promoted a significant increase in the gene expression of several mitochondrial respiratory complexes in the HF fed offspring (NDUF, Cox8B) (Table 5). To assess whether GDM impaired hepatic mitochondrial function, we measured oxygen consumption rate (OCR) from isolated liver mitochondria. Oxidative phosphorylation begins with the transfer of electrons to either complex I or complex II; therefore, we measured OCR through each pathway. For complex I mediated respiration, the state I OCR level (basal respiration, substrate in the absence of ADP) was reduced with GDM exposure in LF fed male mice (Fig. 8C). All other complex I respiration states remained unchanged between the experimental groups (Fig. 8 D-F). Complex II mediated mitochondrial respiration was defined by GDM induced reductions in the HF fed offspring (Fig. 8G-J). GDM exposure significantly reduced basal respiration, state III respiration (maximum respiration when ADP is no longer limiting) as well as spare capacity (the difference between basal and maximal respiration) (Fig. 8G-J). State IV respiration (oligomycin-independent respiration, heat), remained similar between all genotypes (Fig. 8E). Thus, the elevation in the gene expression of respiratory complexes may be a compensatory mechanism in the liver of GDM HF fed offspring to maintain mitochondrial function (Table 5). BBR treatment did not significantly improve hepatic CI or CII mediated OCR for any respiratory state measured.

We also assessed hepatic cardiolipin (CL) content since it is well established that this phospholipid is required for normal mitochondrial function (Mejia & Hatch, 2016). Our analysis revealed that HF diet significantly reduced hepatic CL in both Lean and GDM offspring (Fig. 8B) (Table 6). Consistent with the lack of BBR mediated protection against mitochondrial dysfunction, BBR treatment did not restore hepatic CL content (Fig. 8B). Together these data indicate that in our animal model, BBR treatment of offspring had a limited effect on hepatic metabolic parameters.

**Table 6:**
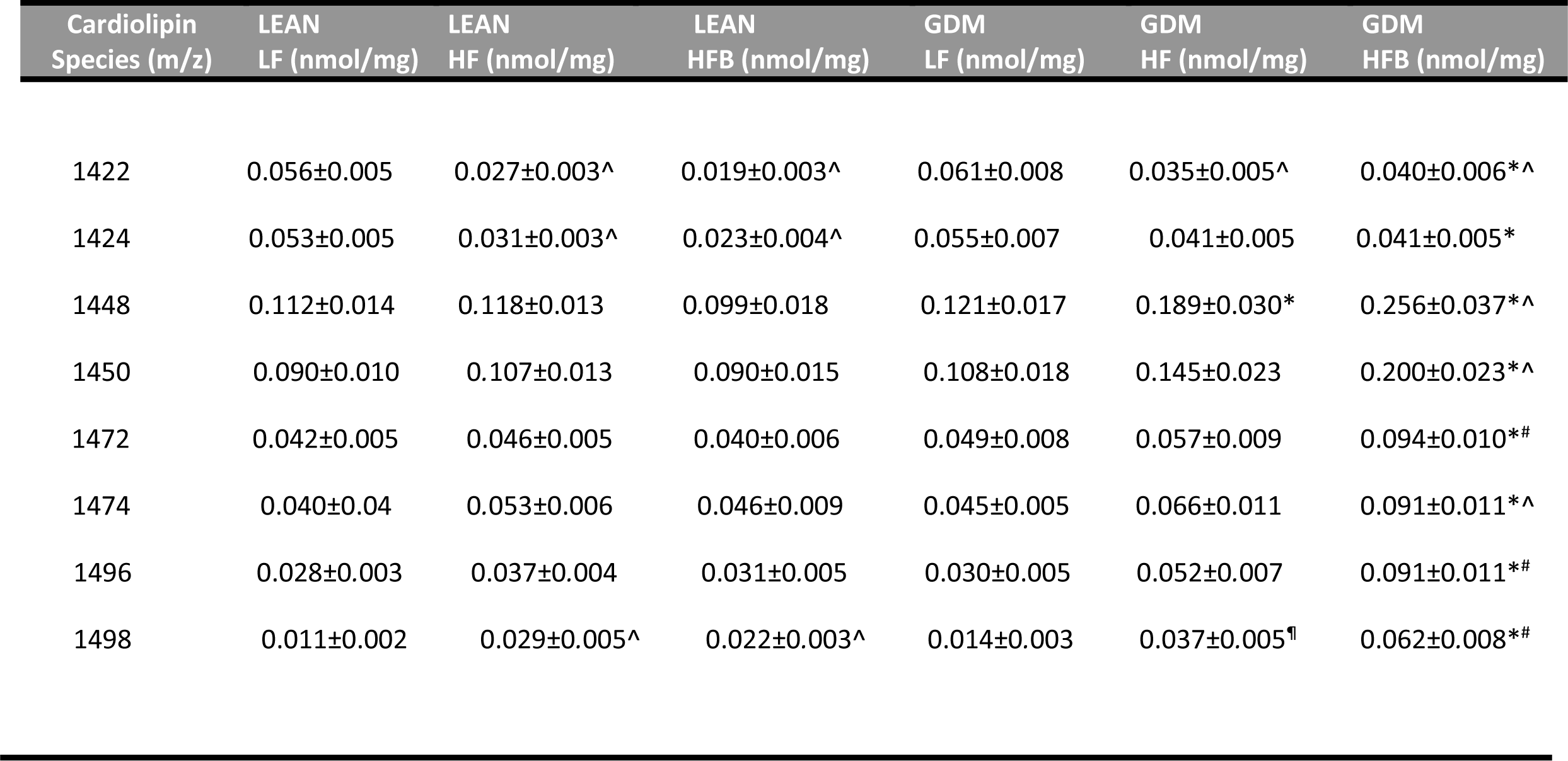
Quantitation of CL species in the liver. Quantitation of molecular species of CL by mass spectrometry from the liver of offspring following 12-weeks of the indicated diet (LF, HF, HFB). Values are means ± SEMs (n=7-10). \**P*<0.05 compared with offspring from lean dams. ¶*P*<0.05 compared to LF and HFB fed offspring from the same gestational condition. ^#^*P*< compared to LF and HF fed offspring from the same gestational condition. ^*P*<0.05 compared to LF fed offspring from the same gestational condition.

### Berberine treatment attenuated cardiac dysfunction due to HF feeding in lean and GDM male offspring

There is accumulating evidence that offspring exposed to maternal diabetes are at greater risk for developing cardiovascular disease (Lee *et al*., 2007; Yu *et al*., 2019). It is also emerging that BBR provides cardioprotection in animal models of heart failure and diabetes (Dong *et al*., 2011; Chang *et al*., 2015b; Chang *et al*., 2016; Dong *et al*., 2018; Feng *et al*., 2019). Therefore, we evaluated the protective effect of BBR on the heart function of GDM exposed male offspring.

We determined that the introduction of a HF diet to the Lean gestational group (HF Lean) induced cardiac dysfunction. Analysis by echocardiography indicated significantly higher isoventricular contraction time (IVCT, 97%) and isoventricular relaxation time (IVRT, 70%) indicating impaired left ventricular systolic and diastolic function respectively in the HF fed animals (Fig. 9A and B). Other variables of systolic function including ejection fraction and fractional shortening were also impaired in the HF group compared to the LF fed controls (LF Lean, Fig. 9C and D). Myocardial dysfunction in the Lean HF group occurred in the absence of any detectable left-ventricular structural remodelling or elevation in arterial blood pressure values (Fig. 9G, Table 7). In the presence of GDM exposure, the HF fed group sustained the pattern of cardiac dysfunction without any additional impairment. However, GDM exposure in the LF fed offspring resulted in significant systolic and diastolic dysfunction with impairments in ejection fraction and fractional shortening (Fig. 9A-F and Table 7).

**Figure 9:**
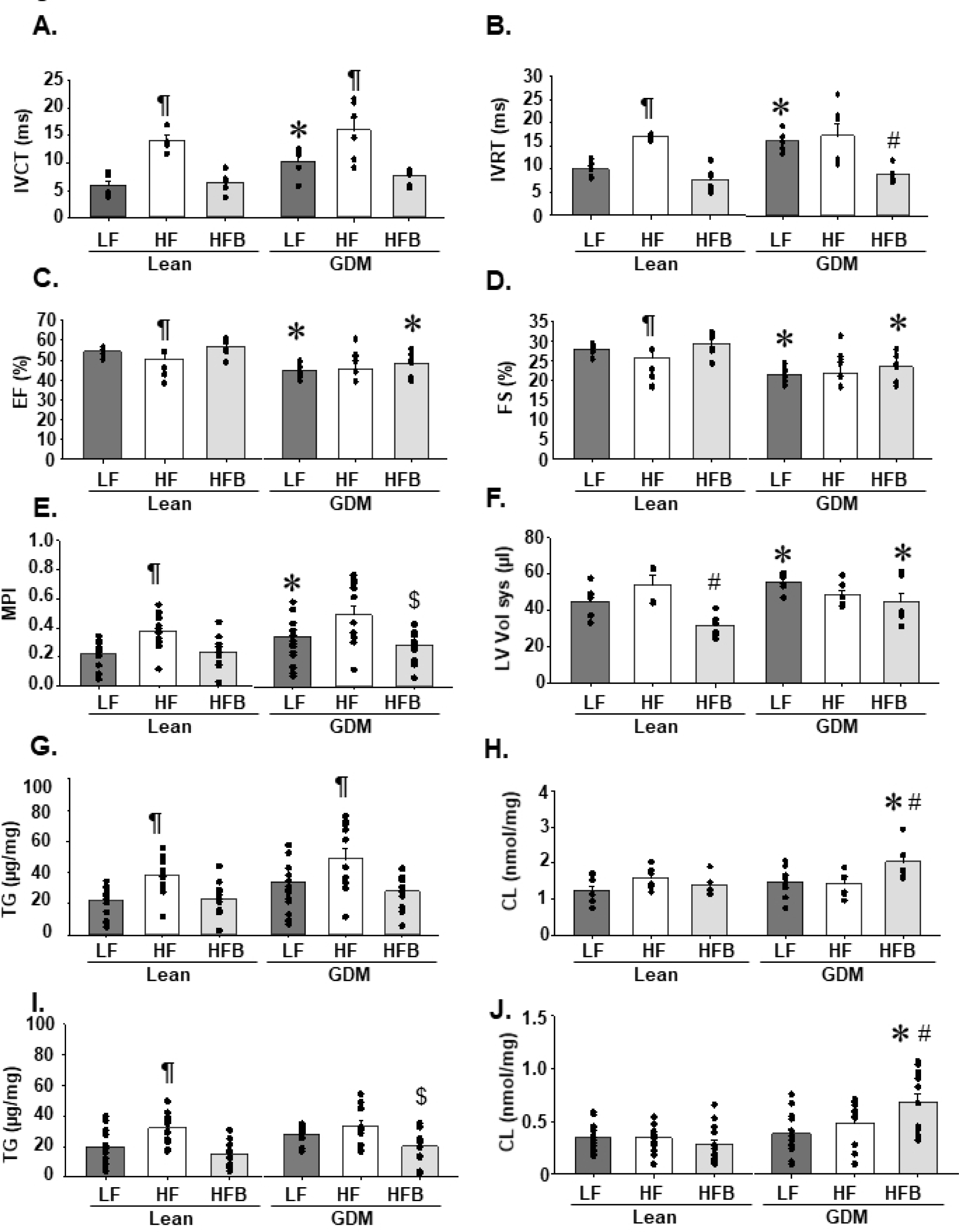
Berberine treatment attenuated cardiac dysfunction due to HF feeding in lean and GDM offspring. Transthoracic echocardiography was performed to determine A, isoventricular contraction time (IVCT), B, isoventricular relaxation time (IVRT), C, ejection fraction (EF), D, fractional shortening (FS), E, myocardial performance index (MPI, isovolumetric contraction time + isovolumetric relaxation time)/ejection time) and F, left ventricular (LV) volumes during systole for male offspring from lean and GDM dams fed the indicated diet for 12 weeks. Quantitation of G, cardiac triglyceride (TG) and H, total CL content. I, Quantitation of skeletal triglyceride (TG) and J, total CL content. All measurements were obtained from male offspring following 12 weeks of the indicated diet (LF, HF, HFB). Values are means ± SEMs. \**P*<0.05 compared with offspring from lean dams. ¶*P*<0.05 compared to LF and HFB fed offspring from the same gestational condition. ^#^*P*< compared to LF and HF fed offspring from the same gestational condition. ^$^*P*<0.05 compared to HF fed offspring from the same gestational condition.

**Table 7:** Echocardiography parameters of the heart. Echocardiography parameters obtained from offspring following 12-weeks of the indicated diet (LF, HF, HFB). Values are means ± SEMs (n=4-22). \**P*<0.05 compared with offspring from lean dams. ¶*P*<0.05 compared to LF and HFB fed offspring from the same gestational condition. ^#^*P*<0.05 compared to LF and HF fed offspring from the same gestational condition. ^*P*<0.05 compared to LF fed offspring from the same gestational condition. d indicates diastolic; s indicates systolic; HW, heart weight; BW, body weight; HR, heart rate; SAP, systolic arterial pressure; DAP, diastolic arterial pressure; MAP, mean arterial pressure; LV, left ventricle; IVRT, isovolumetric relaxation time; MPI, myocardial performance index, was calculated using the equation (isoventricular contraction time + isoventricular relaxation time) / ejection time; LVID, left ventricle internal dimension; LVAW, left ventricle anterior wall; LVPW, left ventricle posterior wall; IVS, interventricular septum.

| Parameter | LEAN<br>LF | LEAN<br>HF | LEAN<br>HFB | GDM<br>LF | GDM<br>HF | GDM<br>HFB |
| --- | --- | --- | --- | --- | --- | --- |
| HW (g) | 0.154±0.005 | 0.160±0.006 | 0.155±0.004 | 0.148±0.003 | 0.145±0.005 | 0.147±0.005 |
| HW/BW (g/g) | 0.0061±0.0002 | 0.0060±0.0002 | 0.0069±0.0003<sup>#</sup> | 0.0052±0.0002<sup>*</sup> | 0.0049±0.0003<sup>*</sup> | 0.0061±0.0003<sup>*#</sup> |
| HR (bpm) | 450±10 | 430±17 | 454±15 | 411±11 | 425±20 | 433±33 |
| SAP (mm Hg) | 81.6±1.9 | 77.5±2.7 | 75.7±1.2 | 75.7±2.2 | 79.1±2.9 | 80.1±1.2 |
| DAP (mm Hg) | 65.7±2.7 | 70.7±2.5 | 67.9±1.8 | 63.2±1.5 | 69.9±2.7 | 67.0±1.1 |
| MAP (mm Hg) | 58±3.0 | 67.5±2.5 | 64.3±2.2 | 57.2±2.0 | 65.7±2.8 | 60.8±1.3 |
| LV d (μl) | 96.1±10.1 | 86.7±5.3 | 68.3±3.2<sup>#</sup> | 98.8±2.7 | 87.2±1.4<sup>¶</sup> | 77.1±4.3<sup>^</sup> |
| LVID s (mm) | 3.31±0.13 | 3.57±0.15 | 2.86±0.10<sup>#</sup> | 3.64±0.06<sup>*</sup> | 3.43±0.08 | 3.31±0.15<sup>*</sup> |
| LVID d (mm) | 4.55±0.20 | 4.37±0.11 | 3.95±0.08<sup>^</sup> | 4.62±0.05 | 4.39±0.03<sup>¶</sup> | 4.16±0.10<sup>#</sup> |
| LVPW s (mm) | 0.96±0.05 | 0.93±0.06 | 0.89±0.05 | 0.88±0.03 | 0.93±0.03 | 0.88±0.06 |
| LVPW d (mm) | 0.68±0.04 | 0.69±0.06 | 0.59±0.03 | 0.64±0.03 | 0.67±0.03 | 0.62±0.04 |
| LVAW s (mm) | 1.19±0.05 | 0.95±0.03<sup>^</sup> | 1.03±0.04<sup>^</sup> | 0.99±0.04<sup>*</sup> | 1.10±0.09 | 0.93±0.05 |
| LVAW d (mm) | 0.75±0.05 | 0.62±0.04 | 0.68±0.03 | 0.71±0.03 | 0.74±0.04 | 0.67±0.04 |
| IVS d (mm) | 1.12±0.06 | 1.03±0.02 | 1.03±0.06 | 1.03±0.02 | 1.04±0.05 | 1.01±0.05 |
| IVS s (mm) | 0.82±0.03 | 0.73±0.04 | 0.71±0.03 | 0.78±0.03 | 0.76±0.05 | 0.76±0.03 |

Interestingly, BBR treatment protected against the development of cardiac dysfunction in both Lean and GDM offspring (Fig. 9A-D). BBR improved the contractility of the left ventricle in the HF fed offspring (Lean and GDM HFB), as reflected by the normalized myocardial performance index (MPI, Fig. 9E). Similar to previous reports, BBR improved both diastolic and systolic function, with IVRT and IVCT restored to normal values (Fig. 9A and B) (Dong *et al*., 2018). However, the protective effects of BBR on ejection fraction and fractional shortening were restricted to the HFB fed Lean gestational group (Fig. 9C and D). We also determined that BBR treatment correlated with reduced left-ventricle systolic and diastolic volume (Fig. 9F and Table 7). Overall BBR treatment ameliorated cardiac dysfunction in both HF fed Lean and GDM exposed offspring with significantly improved markers of left ventricular contractility (IVCT, IVRT, MPI) (Fig. 9A, B and E).

### Berberine treatment attenuated steatosis and mitochondrial dysfunction in the heart of GDM exposed male offspring

Our next objective was to elucidate the mechanism for BBR mediated cardio protection in the GDM exposed offspring. Since, abnormal lipid accumulation is associated with the development of cardiac dysfunction (D’Souza *et al*., 2016), we assessed whether cardiac lipid homeostasis was altered in our animal model. Consistent with the role of BBR as a lipid-lowering agent, we determined that the cardioprotective effect of BBR was mirrored by reduced TG accumulation in heart (Fig. 9G). Specifically, the elevation in TG (∼50%) which occurred in the heart of both the HF fed Lean and GDM exposed offspring was normalized with BBR treatment (Fig. 9G). A similar BBR mediated reduction in TG accumulation was observed in skeletal muscle (Fig. 9I). In the heart, reduced TG levels correlated with increased gene expression of the FAO enzyme ACOX1 (Table 8).

**Table 8:** PCR analysis of the heart. mRNAs were measured from the heart of offspring following 12-weeks of the indicated diet (LF, HF, HFB).. Values are means ±SEMs (n=7). \**P*<0.05 compared with offspring from lean dams. ¶*P*<0.05 compared to LF and HFB fed offspring from the same gestational condition. ^*P*<0.05 compared to LF fed offspring from the same gestational condition. ^$^*P*<0.05 compared to HF fed offspring from the same gestational condition. The abbreviations for the mRNAs are: TAZ, taffazin; PGS, phosphatidylglycerolphosphate synthase; PTPMT, protein tyrosine phosphatase; PGC1α, peroxisome proliferator-activated receptor gamma coactivator 1α; SIRT1, sirtuin 1; CPT1β, carnitine palmitoyltransferase 1; ACOX1, acyl-CoA oxidase; FABP, fatty acid binding protein; NDUF, NADH dehydrogenase oxidoreductase core subunit S3; SDHD, succinate dehydrogenase complex subunit D; COX8β, cytochrome c oxidase subunit 8β; ATP5i, ATP synthase subunit 5i.

| Gene | LEAN<br>LF | LEAN<br>HF | LEAN<br>HFB | GDM<br>LF | GDM<br>HF | GDM<br>HFB |
| --- | --- | --- | --- | --- | --- | --- |
| **CL Synthesis** |  |  |  |  |  |  |
| Tafazzin | 1.00±0.09 | 1.38±0.11<sup>¶</sup> | 0.87±0.05 | 1.16±0.08 | 1.19±0.10 | 1.28±0.05* |
| PGS | 1.00±0.21 | 1.38±0.05<sup>¶</sup> | 0.90±0.10 | 1.13±0.13 | 1.40±0.15 | 1.06±0.08 |
| PTPMTI | 1.00±0.10 | 1.00±0.09 | 0.92±0.04 | 1.00±0.22 | 1.15±16 | 1.10±0.24 |
| **FAO** |  |  |  |  |  |  |
| PGC1α | 1.00±0.14 | 1.12±0.09 | 1.02±0.12 | 1.79±0.21* | 1.62±0.18* | 1.46±0.15* |
| SIRT1 | 1.00±0.15 | 0.91±0.07 | 0.79±0.04 | 1.05±0.22 | 0.82±0.06 | 1.14±0.10 |
| CPT1β | 1.00±0.16 | 1.16±0.19 | 0.83±0.15 | 1.06±0.22 | 1.51±0.17 | 1.52±0.22* |
| ACOX1 | 1.00±0.23 | 0.37±0.10 | 1.45±0.28<sup>\$</sup> | 1.10±0.32 | 1.13±0.28* | 1.40±0.31 |
| **FA Uptake** |  |  |  |  |  |  |
| CD36 | 1.00±0.10 | 0.50±0.06<sup>^</sup> | 0.54±0.09<sup>^</sup> | 1.37±0.14 | 0.65±0.07<sup>¶</sup> | 1.02±0.14* |
| FATBP1 | 1.00±0.14 | 0.52±0.05<sup>^</sup> | 0.50±0.08<sup>^</sup> | 0.63±0.10* | 0.64±0.04 | 1.00±0.19* |
| **ETC** |  |  |  |  |  |  |
| NDUF (I) | 1.00±0.10 | 0.67±0.04<sup>^</sup> | 0.70±0.09<sup>^</sup> | 1.15±0.18 | 0.84±0.08 | 1.01±0.17 |
| SDHD (II) | 1.00±0.20 | 0.78±0.06 | 0.54±0.09 | 0.90±0.24 | 0.61±0.09 | 1.24±0.18*<sup>\$</sup> |
| COX5β (IV) | 1.00±0.08 | 1.27±0.06 | 1.02±0.10 | 1.20±0.14 | 1.30±0.15 | 1.26±0.16 |
| ATP5i (V) | 1.00±0.12 | 1.11±0.09 | 0.96±0.13 | 1.05±0.22 | 1.16±0.12 | 1.15±0.27 |

Based on evidence that BBR protected against HF induced mitochondrial dysfunction in muscle (Gomes *et al*., 2012), we assessed whether a similar protective effect occurred in the heart. For complex I mediated respiration, the state I OCR level (basal respiration) was elevated in response to HF diet in the Lean gestational group (Fig. 10A). However, the amount of spare capacity (difference between basal and maximal) remained unchanged since state III respiration (maximum respiration) was also increased by the HF diet (Lean HF, Fig. 10 B-D). A similar pattern of elevated state I and state III respiration was observed for the GDM LF fed offspring (Fig. 10B-D). However, when GDM exposure was combined with HF feeding, the apparent compensatory increases in state I and III activities were not detected, resulting in a significant decrease in spare capacity (GDM HF, Fig. 10D). Reduction in mitochondrial spare capacity indicates reduced flexibility in response to elevated demands on the heart. State IV respiration (oligomycin-independent respiration, heat) was only elevated in response to GDM exposure in the LF fed offspring (Fig. 10C). There were no detectable differences in cardiac CII mediated OCR between the experimental groups for any of the respiratory states measured.

**Figure 10:**
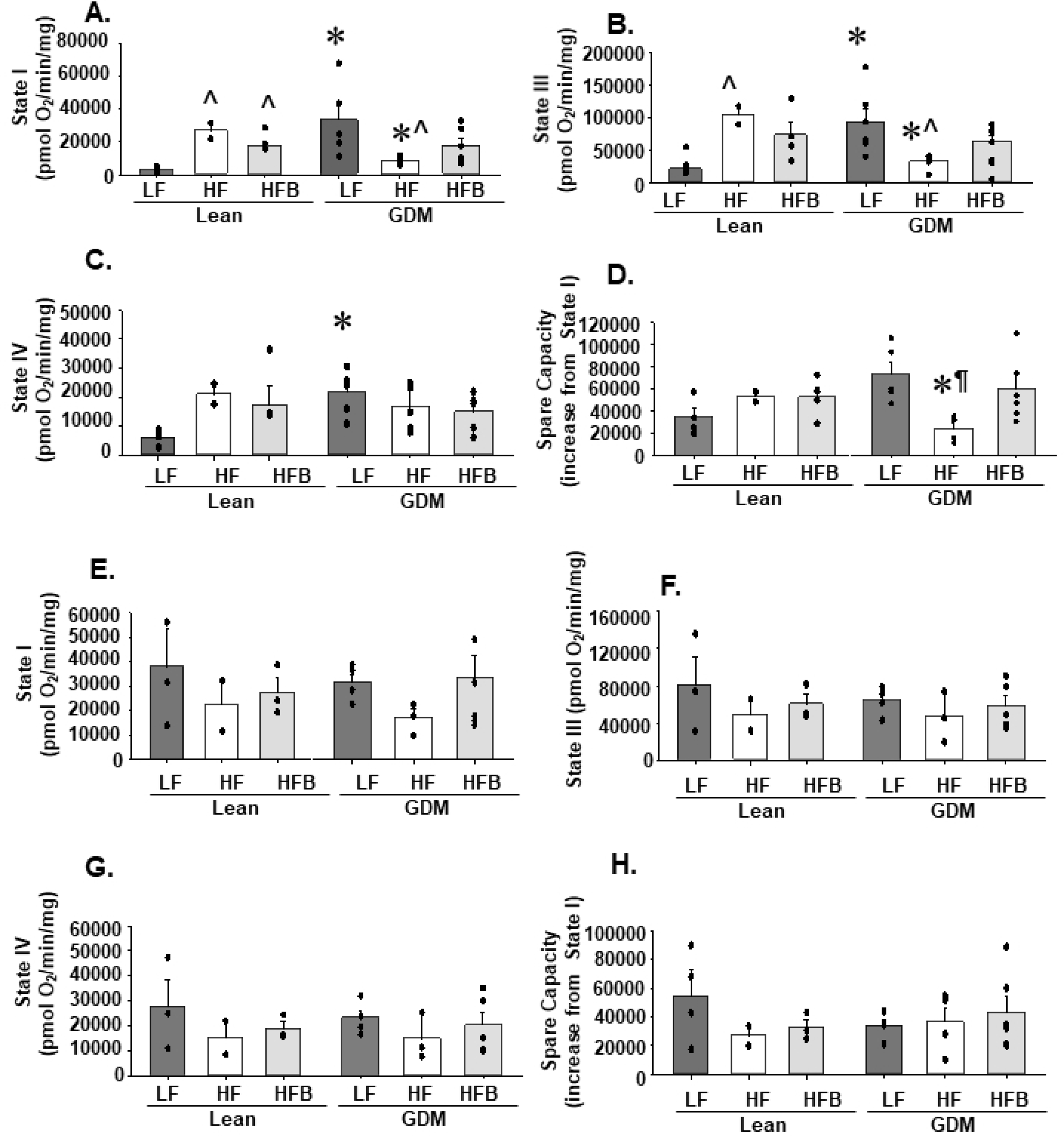
Cardiac complex I spare capacity was improved in GDM offspring by berberine treatment. The oxygen consumption rates (OCR) were measured from cardiac mitochondria isolated from male offspring of Lean and GDM dams fed the indicated diet for 12 weeks. A, Oxygen consumption was measured for Complex I in respiratory state I (basal respiration, substrate in the absence of ADP), B, state III (maximum respiration) and, C state IV (oligomycin (ATP)-independent respiration). D, Spare capacity was calculated using the following formula (state III – state I) / state I *100%). Oxygen consumption was also measured for Complex II in E, respiratory state I (basal respiration, substrate in the absence of ADP), F, state III (maximum respiration, G, state IV (oligomycin (ATP)-independent respiration) and H, spare capacity. All data was normalized to mitochondrial protein content. Values are means ± SEMs. \**P*<0.05 compared with offspring from lean dams. ¶*P*<0.05 compared to LF and HFB fed offspring from the same gestational condition. ^*P*<0.05 compared to LF fed offspring from the same gestational condition.

**Figure 11:**
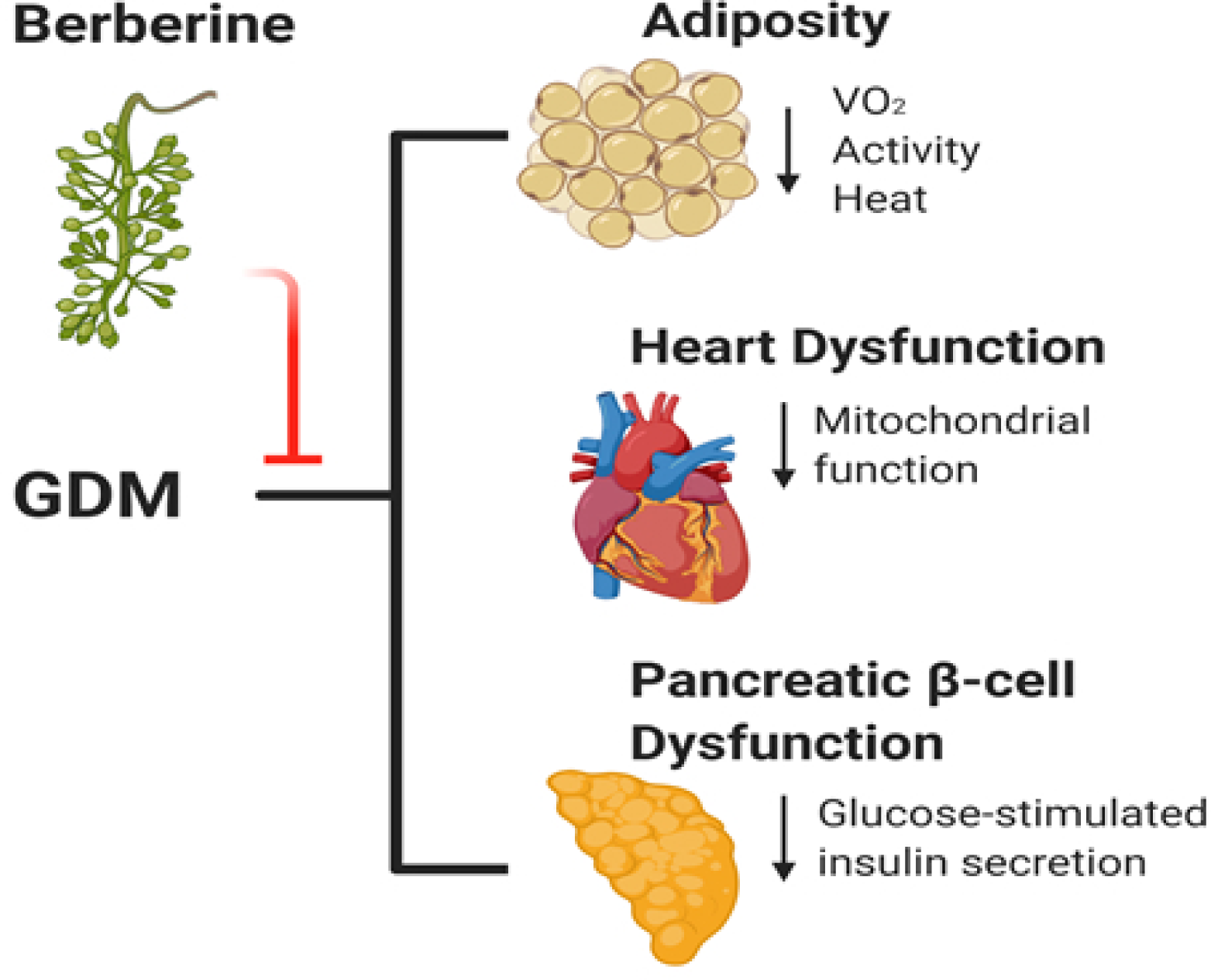
The proposed mechanism for the effects of GDM on HF fed offspring and the preventative effects of BBR. Offspring exposed to GDM had elevated fat mass associated with reduced whole-body metabolism and activity levels. The GDM offspring were additionally susceptible to cardiac contractile, and mitochondrial dysfunction. Finally, glucose-stimulated insulin secretion was reduced from pancreatic islets isolated from GDM exposed offspring. Post-weaning BBR treatment of GDM exposed offspring provided effective treatment from obesity, normalized heart and pancreatic β-cell function.

BBR treatment did not significantly alter CI mediated respiration in the Lean gestational group. However, in the GDM offspring, dietary BBR prevented the HF mediated reductions in state I and state III respiration and thus normalized spare capacity. BBR was also effective at elevating the CL content (∼42%) in the heart and muscle of the GDM exposed groups (Fig. 9H and J). Interestingly, this was due to significant increases (∼32%) in the most abundant and functionally important CL species, tetralinoleoyl cardiolipin (1448) (Table 9 and 10). Alteration in gene expression of CL biosynthetic enzymes was not responsible for the increase in cardiac CL content (Table 8). Together, these results indicate that the BBR mediated protection from cardiac dysfunction in GDM exposed offspring involved its effects on mitochondrial function.

**Table 9:** Quantitation of CL species in the heart. Quantitation of molecular species of CL by mass spectrometry from the heart of offspring following 12-weeks of the indicated diet (LF, HF, HFB). Values are means ± SEMs (n=8). \**P*<0.05 compared with offspring from lean dams. ¶*P*<0.05 compared to LF and HFB fed offspring from the same gestational condition. ^*P*<0.05 compared to LF fed offspring from the same gestational condition.

| Cardiolipin<br>Species (m/z) | LEAN<br>LF (nmol/mg) | LEAN<br>HF (nmol/mg) | LEAN<br>HFB (nmol/mg) | GDM<br>LF (nmol/mg) | GDM<br>HF (nmol/mg) | GDM<br>HFB (nmol/mg) |
| --- | --- | --- | --- | --- | --- | --- |
| 1422 | 0.098±0.010 | 0.036±0.001^ | 0.028±0.004^ | 0.115±0.014 | 0.046±0.008^ | 0.030±0.003^ |
| 1424 | 0.055±0.008 | 0.020±0.004^ | 0.023±0.003^ | 0.057±0.010 | 0.023±0.005^ | 0.030±0.005^ |
| 1448 | 0.569±0.077 | 0.901±0.059^ | 0.718±0.073 | 0.639±0.081 | 0.715±0.075 | 0.943±0.072^* |
| 1450 | 0.180±0.022 | 0.222±0.016^ | 0.232±0.026 | 0.232±0.018 | 0.249±0.037 | 0.283±0.03 |
| 1472 | 0.086±0.03 | 0.100±0.015 | 0.119±0.021 | 0.115±0.016 | 0.113±0.012 | 0.151±0.011 |
| 1474 | 0.050±0.005 | 0.064±0.007 | 0.063±0.006 | 0.052±0.007 | 0.065±0.014 | 0.082±0.011 |
| 1496 | 0.153±0.011 | 0.205±0.024 | 0.233±0.018^ | 0.201±0.030 | 0.228±0.035 | 0.287±0.025 |
| 1498 | 0.044±0.007 | 0.044±0.007 | 0.086±0.020^ | 0.061±0.006 | 0.057±0.01 | 0.107±0.023^ |

**Table 10:**
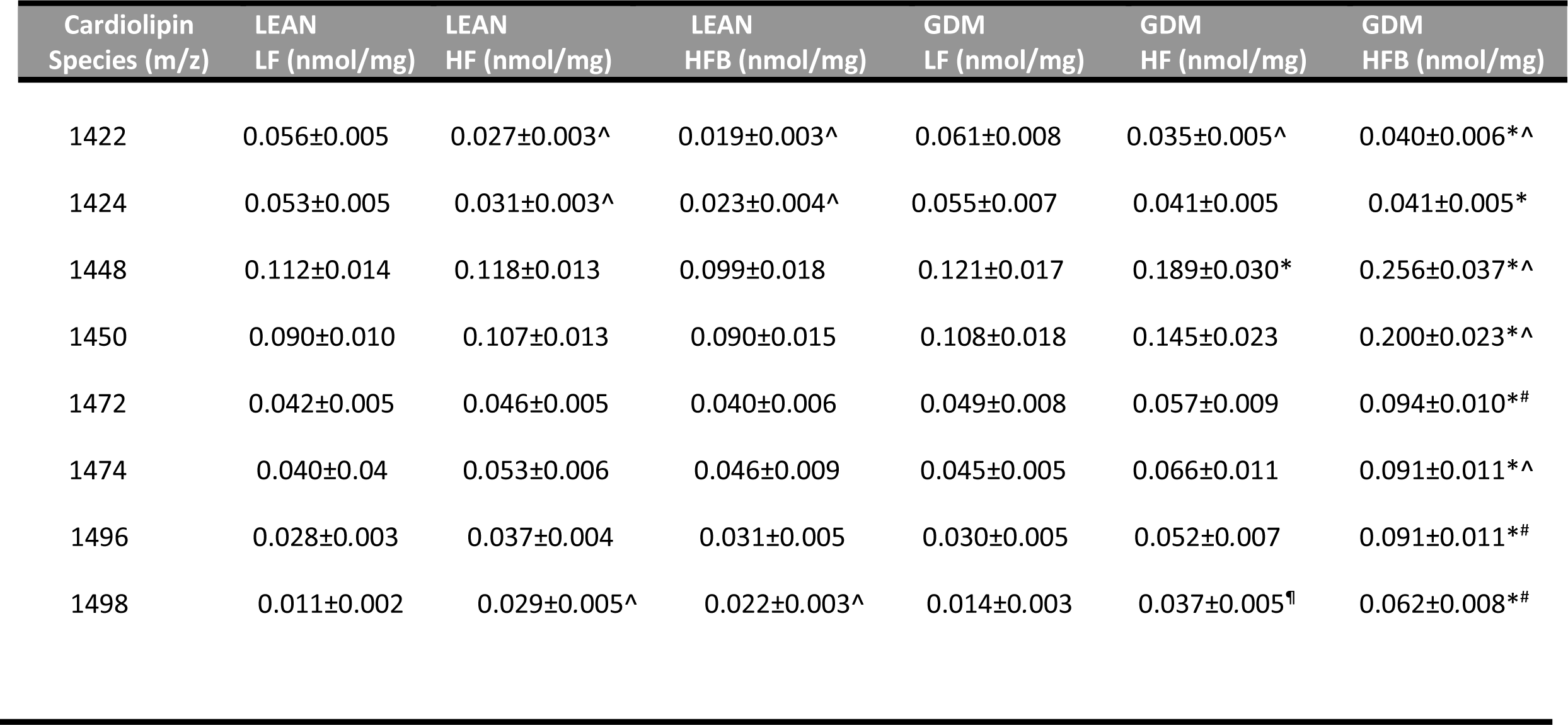
Quantitation of CL in skeletal muscle. Quantitation of molecular species of CL by mass spectrometry from the skeletal muscle of offspring following 12-weeks of the indicated diet (LF, HF, HFB). Values are means ± SEMs (n=13-14). \**P*<0.05 compared with offspring from lean dams. ¶*P*<0.05 compared to LF and HFB fed offspring from the same gestational condition. ^#^*P*< 0.05 compared to LF and HF fed offspring from the same gestational condition. ^*P*<0.05 compared to LF fed offspring from the same gestational condition.

## Discussion

We have determined for the first time that BBR has a protective effect on the adverse metabolic health outcomes of offspring exposed to GDM. We assessed the effectiveness of post-weaning oral BBR supplementation on the metabolic health of high- fat fed offspring exposed to GDM. We established that BBR treatment significantly reduced body weight, fat mass and plasma insulin levels. Importantly, we identified for the first time that dietary BBR reduced adiposity for both sexes in an animal model. We also elucidated a role for BBR in preserving pancreatic β-cell function and heart contractile function in GDM exposed male offspring.

### Dietary BBR supplementation reduced obesity in male and female offspring

Previous reports showed that BBR supplementation reduced the body weight of lean and obese rodents (Lee *et al*., 2006; Zhao *et al*., 2017; Sun *et al*., 2018; Lin *et al*., 2019). BBR also reduced body fat content, with reduced storage of TG in white adipose tissue (Lee *et al*., 2006). Our study supports a BBR mediated reduction in adiposity with decreased % fat mass (Fig 2. G and H), white adipose tissue mass (Fig. 3 A-H) and plasma leptin levels (Table 3 and 4). Little preclinical data is available on the effects of BBR on the metabolic characteristic of females. We have now identified that dietary BBR significantly reduced body weight (∼15-20%), and % fat mass (40-55%) in both male and female offspring. These data are also consistent with the reduced adiposity observed in women with polycystic ovary syndrome treated with BBR (Wei *et al*., 2012).

We identified hypermetabolism in response to BBR for both male and female GDM-exposed offspring. The increase in energy utilization was primarily due to increased heat production (Fig. 4C and D) (Fig. 5 C and D). These results are similar to previous reports of BBR induced thermogenesis in obese db/db male mice (Zhang *et al*., 2014) and HF fed mice (Sun *et al*., 2018; Lin *et al*., 2019). In addition, BBR appears to target white adipose tissue directly to reduce hypertrophy and hyperplasia (Ko *et al*., 2005). A BBR mediated reduction in WAT growth would potentially account for the reduced adiposity in the HFB Lean exposure group which occurred in the absence of elevated energy expenditure (Fig. 4A and B) (Fig. 5A and B).

### Dietary BBR supplementation preserved pancreatic β-cell function

A major finding of our study was demonstrating that the hyperinsulinemia in male GDM offspring was normalized with BBR treatment (Table 3). We reported for the first time chronic dietary BBR protected pancreatic β-cells from elevated insulin secretion under basal/low glucose conditions (Fig. 7C). It was previously demonstrated that insulin secretion was reduced from rat islets pretreated with media containing BBR (Bai *et al*., 2018). Reductions in insulin secretion under low glucose conditions were detected when palmitate was added to the media (Zhou *et al*., 2008). As a result of the palmitate- potentiated insulin secretion, the insulin lowering effect of BBR was detected (Zhou *et al*., 2008). Our results mirror this finding since BBR prevented the GDM induced elevation of insulin secretion in HF-diet fed offspring (Fig. 7C). Notably, the effects of BBR on insulin secretion remains controversial with evidence for both increased and decreased GSIS from rat islets treated (24 h) with BBR (1-10 µM) in the media (Wang *et al*., 2008; Zhou *et al*., 2008). As opposed to the acute use of BBR these preceding studies, our findings suggest that in the setting of GDM exposure, chronic dietary BBR in mice preserved normal GSIS.

### The effects of BBR treatment on hepatic metabolism

The anti-obesity effect of BBR treatment has been correlated with reductions in hepatic steatosis (Zhao *et al*., 2017; Sun *et al*., 2018; Lin *et al*., 2019). In general, significant reductions in hepatic TG accumulation have been achieved with >10% reductions in body weight due to less NEFA released from a smaller mass of adipose tissue (Vilar-Gomez *et al*., 2015). Consistent with this concept we detected lower hepatic TG in the male HFB Lean group which correlated with reduced body weight (15%), fat mass (42%), and plasma NEFA levels (35%) (Table 3). Our results also support prior studies which link BBR mediated hepatic TG reduction with elevated fatty acid oxidation (Table 5) (Brusq *et al*., 2006; Yuan *et al*., 2015; Zhao *et al*., 2017). Overall, the BBR mediated reduction in hepatic TG accumulation in the HFB Lean offspring was relatively small (18%) compared to previous reports (∼20-50%) (Brusq *et al*., 2006; Sun *et al*., 2018) due in part to the absence of HF induced hepatic steatosis.

In the HF GDM offspring, despite significant weight loss (16%) we did not detect an accompanying reduction in hepatic TG levels in response to BBR (Fig. 8A). This may be due to the lack of BBR mediated reductions in plasma lipid levels (Table 3) and unaltered gene expression of hepatic FAO enzymes (Table 5) in GDM exposed offspring. Nevertheless, our findings indicate that BBR treatment consistently improves liver function as indicated by normalization of plasma ALT levels in HF-fed GDM exposed offspring (Table 3).

### Dietary BBR supplementation attenuated GDM-induced mitochondrial and cardiac dysfunction

We have determined for the first time that BBR treatment attenuates the development of cardiac dysfunction in GDM exposed offspring (Fig. 9A-D). Previously, BBR treatment protected against myocardial dysfunction in HF diet and streptozotocin- induced diabetes mice (Chang *et al*., 2015b; Chang *et al*., 2016; Dong *et al*., 2018; Li *et al*., 2018). Consistent with these studies our data shows that BBR significantly reduces both systolic and diastolic left-ventricle dysfunction.

We determined that BBR protects against cardiac dysfunction with an accompanying increase in the heart weight to body weight ratio (Table 7). This is in contrast to previous studies demonstrating that BBR attenuates pathological cardiac hypertrophy (Chang *et al*., 2015b; Dong *et al*., 2018). In our model of GDM, a reduction in heart weight correlated with dysfunction that was reversed with BBR treatment. This suggests that the onset of cardiac hypertrophy may be part of the compensatory mechanism induced by BBR to maintain normal contractile function (Fig. 9A-D and Table 7). Pathways that are known to regulate postnatal cardiac growth are also targeted by BBR (Shiojima *et al*., 2002; Chen *et al*., 2014). The BBR mediated increase in heart mass could not be classified as a traditional physiological hypertrophy since left ventricular wall thickness was within the normal range (Table 7)(Nakamura & Sadoshima, 2018).

One factor that could contribute to contractile dysfunction in these offspring is GDM induced mitochondrial dysfunction. Interestingly, we show that BBR significantly improved mitochondrial respiration in the heart of GDM exposed offspring treated with BBR (Fig. 10A-D). Our results confirm the previously established benefits of dietary BBR on mitochondrial respiration (Gomes *et al*., 2012). Specifically, we show that BBR improved complex 1 function. Notably, complex 1 dysregulation is linked to the initial phases of the failing heart (Rosca *et al*., 2013). In addition, BBR may mediate its cardioprotective effects by increasing CL content (Fig. 9H). CL is essential for maintaining mitochondrial respiration since all energy-transducing enzymes require this lipid. For example, CL binds directly to subunits of complex 1 which promotes global conformational changes and subsequently modulates activity (Jussupow *et al*., 2019). Cell signaling pathways which depend on CL (Dudek & Maack, 2017) may additionally be altered by BBR treatment (Li *et al*., 2019). Together, these studies indicate that dietary BBR effectively attenuates the development of cardiac dysfunction in GDM exposed offspring by improving mitochondrial respiratory function.

## Conclusions

We established that BBR has a protective effect on the health outcomes of offspring exposed to GDM. The HF-fed GDM offspring developed diet-induced obesity, hyperinsulinemia and pancreatic β-cell dysfunction. We determined that BBR treatment significantly reduced body weight and fat mass by promoting hypermetabolism. Furthermore, BBR treatment reversed hyperinsulinemia and preserved pancreatic β-cell function. The HF-fed GDM offspring additionally developed a cardiomyopathy, characterized by both systolic and diastolic dysfunction. BBR treatment attenuated heart contractile dysfunction and mitochondrial respiratory impairment. Our data supports BBR as a potential pharmacotherapeutic approach to improve health outcomes in individuals exposed to GDM.

## Acknowledgments

Supported by grants from the Heart and Stroke Foundation of Canada (GMH), CIHR Grant#144626 (VWD) and the Canadian Foundation for Innovation (VWD). GMH is the Canada Research Chair in Molecular Cardiolipin Metabolism. VWD is the Allen Rouse-Manitoba Medical Services Foundation Basic Scientist.

